# Cross-species dissemination of diverse plasmid lineages and mobile resistance modules shapes carbapenemase co-production in Chilean Enterobacterales

**DOI:** 10.64898/2026.11.22.768343

**Authors:** Nicolás Guerra-Leal, Nahuel Pilquinao, Camilo Berríos-Pastén, Patricio Arros, Matías Gálvez-Silva, Joaquín Acosta, Aaron Ascencio, Erika Valdivia, Rosalba Lagos, Francisco P. Chávez, Ingrid Araya, Rodrigo Iglesias, Pamela Araya, Juan Carlos Hormazábal, Aura Villamil, Andrés E. Marcoleta

## Abstract

Carbapenemase co-production among Enterobacterales is an emerging clinical challenge, yet the relative contributions of clonal expansion, cross-species plasmid dissemination, and movement of resistance modules remain unclear. We integrated national surveillance data with plasmid-resolved hybrid genomes to investigate these processes in Chile. Among 414 isolates carrying at least two carbapenemase families referred during 2018–2024, annual numbers increased markedly alongside broader taxonomic and geographic representation. Thirty isolates were selected to span variation in year, species, region, and carbapenemase profile. These represented 11 species from six genera and contained 149 complete plasmids, 57 of which carried carbapenemase genes. Of 63 carbapenemase gene copies, 62 were plasmid-borne, and 55 of the 57 carbapenemase-carrying plasmids were predicted to be conjugative or mobilizable. Complementary plasmid typing, whole-sequence clustering, structural comparisons, and pan-plasmidome analysis identified three recurrent lineages: IncX3/*bla*_NDM-7_, IncN/*bla*_KPC-2_, and IncHI2A/*bla*_VIM-1_. Their occurrence across multiple species and sequence types supported cross-species dissemination, while numerous minor groups and singletons revealed a broader repertoire of plasmid architectures and accessory genes. Related backbones carried variable resistance regions, and equivalent class 1 integrons and transposon-associated platforms occurred in otherwise unrelated plasmids, indicating modular redistribution of carbapenemase genes. In selected isolates, co-resident plasmids created resistance redundancy, brought recurrent carbapenemase plasmid lineages together, or coupled resistance and virulence within the same cell. Together, these findings indicate that carbapenemase co-production results from the combined contributions of clonal dissemination, cross-species plasmid transfer, and the horizontal movement of mobile resistance modules. Complete plasmid resolution is essential to distinguish these processes and track their dissemination.

## INTRODUCTION

Antimicrobial resistance (AMR) is a major and growing cause of morbidity and mortality worldwide. In 2021, bacterial AMR was associated with an estimated 4.71 million deaths, including 1.14 million attributable deaths, and its burden is projected to increase substantially by 2050 (Naghavi et al., 2024). Carbapenem-resistant Enterobacterales (CRE) are particularly concerning because carbapenems are often among the last reliable treatments for severe infections caused by multidrug-resistant Gram-negative bacteria. Accordingly, the 2024 World Health Organization Bacterial Priority Pathogens List classifies CRE as critical-priority pathogens, with carbapenem-resistant *Klebsiella pneumoniae* receiving the highest score among individually assessed pathogens (Sati et al., 2025; WHO, 2024).

Although carbapenem resistance can result from reduced permeability combined with extended-spectrum β-lactamases or AmpC cephalosporinases, its most epidemiologically significant mechanism is carbapenemase acquisition (Bonomo et al., 2018; Hammoudi Halat and Ayoub Moubareck, 2020). The principal families comprise class A KPC enzymes, class B metallo-β-lactamases such as NDM, VIM, and IMP, and class D OXA-48-like enzymes. Most carbapenemase-producing Enterobacterales carry a single family, but isolates co-producing two or more are increasingly reported, including in Latin America and the Caribbean (Pan American Health Organization/World Health Organization, 2021; Yuan et al., 2024). Combinations of serine carbapenemases and metallo-β-lactamases are clinically important because their complementary hydrolytic and inhibitor-susceptibility profiles can compromise enzyme-directed therapeutic strategies (Tamma et al., 2024). Such isolates may therefore serve as sentinels of advanced resistance evolution.

The epidemiology of carbapenemase co-production depends not only on bacterial clones but also on the mobile platforms carrying the corresponding genes. Carbapenemase genes are frequently embedded in insertion sequences, transposons, and integrons located on conjugative or mobilizable plasmids (Partridge et al., 2018). Recurrent associations reported globally include KPC with IncF and IncN plasmids, NDM with IncX3 plasmids, and OXA-48-like enzymes with IncL plasmids, although the same gene can occur in multiple backbones and genetic contexts (Rozwandowicz et al., 2018). Plasmids may also carry determinants of resistance to other antimicrobials, metals, and biocides and, less frequently, virulence-associated loci, creating opportunities for co-selection and resistance–virulence convergence (Gálvez-Silva et al., 2024; Pal et al., 2015).

Genomic epidemiology has greatly expanded the description of carbapenemase-producing Enterobacterales, but many studies rely on short-read draft assemblies (Arredondo-alonso et al., 2017). These datasets are well suited to lineage and resistance-gene surveillance. However, they often cannot unambiguously assign determinants to individual plasmids, resolve nested integrons and transposons, or distinguish dissemination of a conserved plasmid backbone from repeated movement of the same resistance module. Studies based on complete, plasmid-resolved genomes therefore remain comparatively uncommon (Arredondo-alonso et al., 2017; Bouras et al., 2024). This limitation is particularly relevant in underrepresented regions of the Global South. A regional review covering publications through 2019 identified carbapenemase-associated mobile elements in only seven South American countries, with the available evidence centered largely on particular clones, incompatibility groups, or transposable elements (Reyes et al., 2020). More recent hybrid-sequencing studies from Brazil and Argentina provided valuable plasmid-level resolution but examined individual outbreaks or small sets of specific carbapenemase-bearing plasmids (Boralli et al., 2023; González-Espinosa et al., 2024).

Chile provides an informative setting in which to examine these processes. Following the first detection of KPC-producing *K. pneumoniae* in 2012, national surveillance documented a progressive increase in carbapenemase-producing Enterobacterales. Between 2017 and 2024, 72 healthcare-associated outbreaks involving 945 cases were recorded in public hospitals. From 2021 onwards, these outbreaks encompassed an increasingly diverse range of species and carbapenemase profiles (Lara et al., 2026). Consistently, a multicenter study of invasive CRE isolates collected during 2019–2022 identified isolates co-producing KPC and NDM in four species from several hospitals (Quesille-Villalobos et al., 2025). However, species identification and carbapenemase profiling alone cannot determine whether this diversification reflects clonal expansion, independent acquisition, or cross-species dissemination of shared mobile platforms.

Available Chilean genomic studies illustrate this complexity. A multispecies outbreak in 2015 was associated with cross-species dissemination of a broad-host-range IncN plasmid carrying *bla*_KPC-2_ within NTEKPC-IIe (Wozniak et al., 2021). Related IncN/KPC-2 plasmids were subsequently identified in *K. pneumoniae* and in invasive *Escherichia coli* and *Klebsiella oxytoca*, whereas comparatively conserved IncX3/*bla*_NDM-7_ plasmids occurred across several species (Quesille-Villalobos et al., 2025; Veloso et al., 2023). Additional studies documented acquisition of carbapenemase plasmids by hypervirulent *K. pneumoniae* ST23/KL1, including an IncN plasmid carrying both *bla*_KPC-2_ and *bla*_VIM-1_ (Gálvez-Silva et al., 2024; Morales-León et al., 2023). Together, these studies revealed recurrent plasmid lineages and exceptional resistance–virulence convergence events but remained restricted to particular species, outbreaks, clinical networks, or enzyme combinations. The contributions of prevalent plasmid lineages, less frequent architectures, and transferable resistance modules to carbapenemase co-production across Chilean Enterobacterales therefore remain unresolved.

Here, we examined the temporal, taxonomic, and geographic distribution of Enterobacterales carrying at least two carbapenemase families among isolates referred to the Chilean Public Health Institute between 2018 and 2024 as part of the national surveillance program for carbapenem-resistant Enterobacterales. We then used antimicrobial susceptibility testing and hybrid genome sequencing to characterize a subset of 30 isolates spanning bacterial hosts, carbapenemase combinations, years, regions, clonal backgrounds, and resistance phenotypes. By integrating phylogenomics, resistome and virulence profiling, plasmidome analysis, and reconstruction of integrons and transposable platforms, we examined the roles of clonal expansion, cross-species plasmid dissemination, and modular gene transfer in carbapenemase co-production.

## MATERIALS AND METHODS

### Surveillance dataset and isolate selection

Records of carbapenem-resistant Enterobacterales submitted to the Chilean Public Health Institute (Instituto de Salud Pública de Chile; ISP) through routine national laboratory surveillance between 2018 and 2024 were reviewed to identify isolates carrying at least two carbapenemase families. Carbapenemase families were confirmed at the ISP using family-specific PCR and immunochromatographic assays and grouped as KPC, NDM, VIM, or OXA-48-like. Records were consolidated using the unique ISP isolate identifier. The final dataset comprised 414 unique isolate records: 413 carrying two carbapenemase families and one carrying KPC, NDM, and VIM. Preliminary genus-and species-level identifications were obtained at the ISP by matrix-assisted laser desorption/ionization time-of-flight mass spectrometry (MALDI-TOF MS). Additionally, isolation date, specimen source, and region of origin were obtained from the available surveillance metadata.

Thirty available isolates were selected to cover the widest possible range of carbapenemase-family combinations, preliminary taxonomic assignments, geographic regions, and years represented in the surveillance collection. The panel spanned 2018-2024 and, after genome-based taxonomic resolution, comprised 11 species from six genera.

### Culture conditions and antimicrobial susceptibility testing

Antimicrobial susceptibility testing was performed at the ISP according to the standardized procedures of the national reference laboratory, using broth microdilution and Etest gradient diffusion as appropriate (Veloso et al., 2023). The analysis was restricted to ceftazidime, cefepime, cefotaxime, meropenem, imipenem, ertapenem, colistin, and tigecycline, for which results were consistently available across the 30-isolate panel. MICs were retrospectively interpreted according to the Clinical and Laboratory Standards Institute M100 standard, 33rd edition, for agents with applicable CLSI breakpoints. Tigecycline MICs were reported descriptively without categorical interpretation. Isolate-level MICs and categorical interpretations are provided in Table S2.

### Genome sequencing, assembly, and general annotation

Genomic DNA was extracted using the GeneJET Genomic DNA Purification Kit (Thermo Fisher Scientific, USA) and quantified with a Qubit fluorometer (Invitrogen, USA). Illumina TruSeq Nano libraries were sequenced as 100-bp paired-end reads on a HiSeq 4000 platform by SeqCenter (USA). Oxford Nanopore libraries were prepared in-house using the SQK-RBK004 Rapid Barcoding Kit and sequenced with FLO-MIN106 flow cells on a MinION device. Signals were base-called with Guppy v6.5.7+ca6d6af using the high-accuracy dna_r9.4.1_450bps_sup.cfg model.

Illumina reads were processed with fastp v0.24.0 (Chen et al., 2018) by trimming 21 nucleotides from the 5′ end and five nucleotides from the 3′ end of both paired-end reads (-f 21-t 5-F 21-T 5). Read quality was assessed with FastQC v0.12.1 and NanoPlot v1.42.0 (Andrews, 2010; De Coster and Rademakers, 2023). Long reads were filtered with Filtlong v0.2.1 (https://github.com/rrwick/filtlong), retaining reads ≥1,000 bp, the highest-quality 90% of the input, and up to 500 Mb of sequence (--min_length 1000 --keep_percent 90 -- target_bases 500000000). Hybrid assemblies were generated with Hybracter v0.7.4 (Bouras et al., 2024) using the --auto and --no_medaka options; all remaining parameters were left at their default values. Assembly graphs were inspected with Bandage v0.9.0 (Wick et al., 2015) to verify replicon structure and circularization. Completeness and contamination were estimated with CheckM2 v1.0.2 (Chklovski et al., 2023) using default parameters, and assembly statistics were calculated from the final assemblies with QUAST v5.3.0 (Gurevich et al., 2013) using default parameters. Assemblies were annotated with Bakta v1.9.4 (Schwengers et al., 2021) and the full database v5.1 (19 January 2024), providing the corresponding genus and, when recognized by Bakta, the species assignment through the -- genus and --species options.

### Genome-based taxonomic identification, sequence typing, and phylogenomic analysis

Genome-based taxonomic assignments were obtained with GTDB-Tk v2.4.0 and Genome Taxonomy Database release 220 (Chaumeil et al., 2022). Assignments were independently evaluated by average nucleotide identity against representative and closely related genomes using skani v0.2.2, with 95% ANI as the general species-level threshold (Shaw and Yu, 2023). Discrepancies between the preliminary MALDI-TOF MS identification and the genome-based assignment were resolved using the combined GTDB-Tk, ANI, and phylogenomic evidence. Multilocus sequence types were assigned using Pathogenwatch (accessed in December 2024). Because MLST schemes are species-specific, recurrent and singleton sequence types were evaluated within each species.

To place the resistome of the study isolates in a broader phylogenetic context, species-specific public-genome datasets were assembled in December 2024. For all species except *K. pneumoniae*, all assemblies available in NCBI GenBank were retrieved using NCBI Datasets v16.34.1. Because few *Leclercia tamurae* genomes were available, the initial retrieval was expanded to include all genomes assigned to the genus *Leclercia*. For *K. pneumoniae*, the Chilean isolates were first examined using Pathogenwatch to define their phylogenetic context. Public genomes representing the STs identified among the Chilean isolates were then selected to provide the most closely related references available for each ST, yielding a comparison set of 120 genomes. Across all datasets, assemblies flagged as atypical or derived from metagenomes were excluded, and only genomes with CheckM2-estimated completeness >95% and contamination <5% were retained. Taxonomic assignments were subsequently verified with GTDB-Tk. Species-specific datasets retained only genomes assigned to the corresponding species, whereas the *Leclercia* dataset retained all genomes assigned to that genus to provide genus-wide context for the *L. tamurae* study isolate.

A pangenome was constructed separately for each species with Panaroo v1.5.0 using strict cleaning and a 95% core-gene threshold (--clean-mode strict --core_threshold 0.95) (Tonkin-Hill et al., 2020). Concatenated core-gene nucleotide alignments were used to infer maximum-likelihood trees with IQ-TREE v2.3.6 under the GTR+G model and 1,000 ultrafast bootstrap replicates (-m GTR+G-B 1000) (Minh et al., 2020). Trees were mid-point rooted, annotated with isolate metadata, and visualized in iTOL v7 (Letunic and Bork, 2021). Phylogenetic position, sequence type, carbapenemase profile, and relevant resistance or virulence features were examined jointly.

### Resistome profiling

Antimicrobial resistance genes and selected stress-response determinants were identified with AMRFinderPlus v4.0.19 (Feldgarden et al., 2021) and database v2024-10-22.1 using the annotated coding sequences and nucleotide assemblies with default parameters. Determinants were classified by predicted association with antimicrobial resistance, reduced susceptibility to disinfectants or biocides, or metal tolerance. Carbapenemases and other β-lactamases were classified by family and allele based on the beta-lactamase database BLDB (Naas et al., 2017). Copy numbers were calculated from non-overlapping matches on each complete replicon, and location was assigned as chromosomal or plasmid-borne. This approach was also used to examine recurrent associations, including the *bla*_NDM_-*ble* module and co-occurrence of *sul1* and *qacE*Δ*1*.

### Plasmid typing, mobility prediction, and sequence-based clustering

All complete extrachromosomal replicons were analyzed with MOB-suite v3.1.9 (Robertson and Nash, 2018). The mob_typer module was used to identify replicon and incompatibility-group markers, relaxase types, mating-pair formation systems, predicted origins of transfer, host-range assignments, and primary and secondary MOB-cluster identifiers. Plasmids were classified as conjugative, mobilizable, or nonmobilizable according to the MOB-suite output.

Plasmids with identical primary and secondary MOB-cluster assignments were grouped and designated pA-iB, where A denotes the number of plasmids and B is a sequential index assigned after ordering groups from largest to smallest. Thus, p19-i1 and p16-i2 contained 19 and 16 plasmids and represented the first-and second-largest groups, respectively. Equal-sized groups received consecutive indices. Plasmids lacking either MOB-cluster assignment were classified as untyped. Inc groups were analyzed independently because shared Inc markers did not necessarily correspond to the same MOB-cluster pair.

To complement this reference-dependent classification, whole-plasmid sequence profiles were generated for all 149 Chilean plasmids using MGEcluster v1.1.0 with k = 10 (Arredondo-Alonso et al., 2023). Because these profiles are high-dimensional, t-distributed stochastic neighbor embedding was used to produce a two-dimensional representation emphasizing local sequence neighborhoods (t-SNE; perplexity 42, random state 42). HDBSCAN, a hierarchical density-based clustering algorithm (McInnes et al., 2017), was then applied to the t-SNE coordinates with a minimum cluster size of 16. This approach identifies stable regions of high point density without requiring the number of groups to be specified in advance, accommodates groups of different shapes, and retains plasmids lacking a stable density-based association as ungrouped. Membership probabilities provided support for individual assignments. This workflow allowed us to assess whether the MOB-defined groups were also recovered directly from whole-plasmid sequence composition.

For global comparison, the 57 Chilean carbapenemase-positive plasmids were first used to define the target primary-secondary MOB-cluster pairs. Complete plasmids from PLSDB release 2024_05_31_v2 (Galata et al., 2019) were typed with the same MOB-suite workflow and retained only when both MOB-cluster identifiers matched one of these pairs. After defining the target pairs, all Chilean plasmids assigned to them were included in the combined analysis. This added ten carbapenemase-negative members of the same MOB-defined groups, yielding 67 Chilean plasmids. The 1,182 retained PLSDB plasmids and 67 Chilean plasmids were analyzed jointly with MGEcluster using k=10, t-SNE perplexity 42, random state 42, and a minimum HDBSCAN cluster size of 11.

### Sample-based rarefaction analysis

Sampling completeness of the plasmid repertoire was evaluated using sample-based incidence rarefaction. The 30 bacterial isolates were treated as sampling units, and each unique pair of primary and secondary MOB-cluster assignments was considered an operational plasmid lineage. All plasmids were included regardless of carbapenemase carriage, whereas two plasmids lacking either MOB-cluster assignment were excluded. When multiple plasmids assigned to the same lineage occurred within one isolate, the lineage was counted only once, producing a binary isolate-by-lineage incidence matrix. Analytical rarefaction and short-range extrapolation to 60 isolates, twice the observed number of sampling units, were calculated following the sample-based framework of (Colwell et al., 2012) and (Chao et al., 2014). Asymptotic lineage richness was estimated using the bias-corrected Chao2 estimator: SChao2 = Sobs + [(T − 1)/T] × Q1(Q1 − 1)/[2(Q2 + 1)], where T is the number of isolates and Q1 and Q2 are the numbers of lineages detected in exactly one and two isolates, respectively. Incidence-based sample coverage was estimated from the same frequency distribution. Confidence intervals were obtained using 1,000 bootstrap replicates following the procedure described previously (Chao et al., 2014).

### Pan-plasmidome and structural comparison of major plasmid clusters

Gene-content analyses were performed separately for the three predominant plasmid clusters, p19-i1, p16-i2, and p10-i3. Plasmid-specific GFF3 annotations generated with Bakta v1.9.4 and the full database v5.1 (19 January 2024) were analyzed with Panaroo v1.5.0 using --clean-mode sensitive and --remove-invalid-genes. All other parameters were left at their default values. Gene presence-absence matrices were used to quantify each repertoire and classify genes using the default frequency thresholds: core (99–100%), soft core (95–<99%), shell (15–<95%), and cloud (<15%). Matrices were integrated with plasmid length, bacterial host, year, mobility, replicon type, carbapenemase content, and other resistance determinants.

Complete circular sequences and their annotations were inspected to identify differences in backbone and accessory regions. Circular comparisons were generated with pyCirclize v1.10.1 (https://github.com/moshi4/pyCirclize), displaying coding sequences, resistance and virulence-associated genes, insertion sequences, integrons, and other relevant mobile elements. Panaroo gene-content patterns and inspection of the complete plasmid structures were interpreted jointly: conservation of backbone content and order supported a coherent lineage, while accessory regions were examined for insertions, deletions, duplications, and rearrangements. Because Panaroo captures gene-family presence but does not resolve the structural organization or copy number of repeated regions, duplications of carbapenemase genes, *sul1*, and *qacE*Δ*1* were verified directly in the complete plasmid assemblies and corresponding annotations.

### Integrons and genetic environments of carbapenemase genes

Integrons were identified in complete chromosomal and plasmid sequences using the Docker implementation of IntegronFinder v2.0.5 with default parameters (Néron et al., 2022). Predictions were classified according to the presence and organization of an integron integrase, attC sites, and associated gene cassettes. AMRFinderPlus hits were mapped to integrons by genomic coordinates. Carbapenemase genes were considered integron-associated only when located within predicted boundaries or a manually validated cassette array. Integrase sequences were inspected to identify class 1 integrons carrying *intI1* and distinguish them from other classes or incomplete integron-like structures.

Insertion sequences and transposon-associated regions surrounding each carbapenemase copy were identified with ISEScan v1.7.2 using default parameters (Xie and Tang, 2017) and examined together with Bakta and AMRFinderPlus annotations. Flanking regions from the corresponding complete replicon were inspected for insertion-sequence genes, transposases, resolvases, and inverted or direct repeats. Protein homology was evaluated with BLASTp through Biopython (Cock et al., 2009), retaining alignments of at least 100 amino acids and 98% identity. The corresponding nucleotide regions were aligned with MAFFT using the --auto option (Katoh and Standley, 2013). Equivalent platforms in different backbones and repeated copies of *bla*_NDM-7_, *bla*_VIM-2_, *sul1*, and *qacE*Δ*1* were verified directly from complete, long-read-resolved assemblies.

## RESULTS AND DISCUSSION

### Dual-carbapenemase-producing Enterobacterales increased and diversified across Chile between 2018 and 2024

Through Chile’s national AMR surveillance, 414 unique Enterobacterales isolates carrying at least two carbapenemase families were identified between 2018 and 2024 (Fig. 1; Table S1). The earliest such isolate was recorded in 2018, with none detected in previous years. Annual referrals increased from one isolate in 2018 and 2019 to 21 in 2020 and 160 in 2024, representing a 7.6-fold increase since 2020, with only a modest decline in 2022 (Fig. 1A). The sustained rise in referrals, particularly during 2023–2024, indicates a marked expansion of multiple-carbapenemase-producing Enterobacterales within Chile’s national surveillance network.

**Figure 1.**
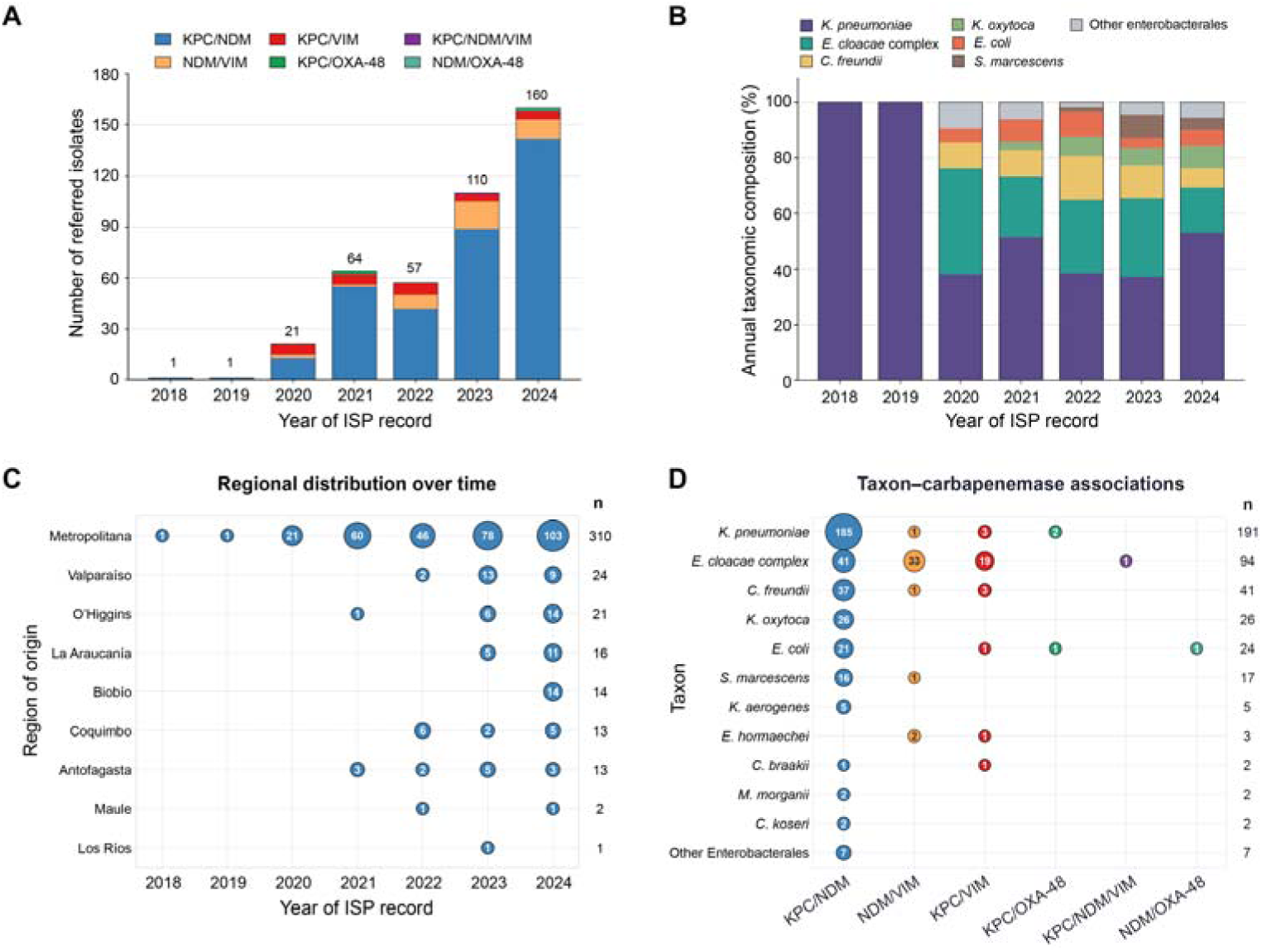
Temporal, taxonomic, and geographic diversification of Enterobacterales carrying at least two carbapenemase families in Chile, 2018–2024. (A) Annual number of isolates received by the Chilean Public Health Institute (Instituto de Salud Pública de Chile; ISP), stratified by detected carbapenemase combination. Numbers above the bars indicate annual totals. (B) Relative annual taxonomic composition of the isolates. The six most frequent taxa are shown separately, and the remaining taxa are grouped as “Other Enterobacterales.” (C) Geographic distribution of isolates according to region of origin and year. Bubble area is proportional to the number of isolates, numbers within bubbles indicate exact counts, and values on the right indicate regional totals. (D) Distribution of carbapenemase combinations across taxa. Taxa represented by at least two isolates are shown separately, whereas taxa represented by a single isolate are grouped as “Other Enterobacterales.” Bubble area is proportional to the number of isolates, numbers within bubbles indicate exact counts, and values on the right indicate taxon totals. The dataset comprised 414 ISP isolate records: 413 carrying two carbapenemase families and one *E. cloacae* complex isolate carrying KPC, NDM, and VIM. Years correspond to the year of the ISP record identifier. KPC, *K. pneumoniae* carbapenemase; NDM, New Delhi metallo-β-lactamase; VIM, Verona integron-encoded metallo-β-lactamase; OXA-48, OXA-48-like carbapenemase.

This numerical expansion was accompanied by a broader range of bacterial hosts. Whereas the isolates recorded in 2018 and 2019 were exclusively *K. pneumoniae*, six taxonomic groups were represented in 2020, 11 in 2023, and 14 in 2024 (Fig. 1B). Overall, the collection comprised 15 named species or species complexes from ten genera, together with three isolates identified only to the genus level. The most frequent taxa were *K. pneumoniae* (191/414, 46.1%), the *Enterobacter cloacae* complex (94/414, 22.7%), *Citrobacter freundii* (41/414, 9.9%), *Klebsiella oxytoca* (26/414, 6.3%), *Escherichia coli* (24/414, 5.8%), and *Serratia marcescens* (17/414, 4.1%), which together accounted for 94.9% of the records. Although the larger number of isolates examined in later years likely contributed to the observed richness, the distribution of multiple-carbapenemase profiles across both common and less frequently reported Enterobacterales indicates that the phenomenon was no longer confined to *K. pneumoniae*.

KPC/NDM was the predominant combination, accounting for 343 isolates (82.9%) and occurring in 17 of the 18 taxonomic categories (Fig. 1D). NDM/VIM and KPC/VIM were detected in 38 (9.2%) and 28 (6.8%) isolates, respectively, whereas KPC/OXA-48-like and NDM/OXA-48-like occurred in three (0.7%) and one (0.2%) isolates. One *E. cloacae* complex isolate carried KPC, NDM, and VIM. Carbapenemase profiles differed substantially between the principal hosts. KPC/NDM accounted for 185 of 191 *K. pneumoniae* isolates (96.9%), whereas the *E. cloacae* complex comprised KPC/NDM (41/94, 43.6%), NDM/VIM (33/94, 35.1%), KPC/VIM (19/94, 20.2%), and KPC/NDM/VIM (1/94, 1.1%). Thus, 53 of 94 *E. cloacae* complex isolates (56.4%) carried a VIM-containing profile. Temporal diversification was also evident: KPC/VIM and NDM/VIM were first recorded in 2020, KPC/OXA-48-like in 2021, KPC/NDM/VIM in 2023, and NDM/OXA-48-like in 2024. These host-specific distributions are inconsistent with expansion of a single carbapenemase combination through a uniform bacterial population.

Geographic representation also broadened over time (Fig. 1C). All records from 2018–2020 originated in the Metropolitan Region, but the cumulative number of regions represented increased from three by the end of 2021 to nine by the end of 2024. Overall, 310 isolates (74.9%) came from the Metropolitan Region and 104 (25.1%) from elsewhere in the country. The non-metropolitan proportion increased from 6.2% in 2021 to 35.6% in 2024, and KPC/NDM was detected in eight of the nine regions represented during the study. The broader geographic coverage highlights the expanding national reach of surveillance, with regional counts reflecting patterns of laboratory referral and diagnostic activity.

These findings are consistent with complementary evidence of increasing diversification in Chile. Lara et al. (2026) (Lara et al., 2026) documented 72 healthcare-associated outbreaks involving 945 cases between 2017 and 2024, with KPC/NDM co-production reported in outbreak surveillance from 2021 onward and cases distributed across 11 of the country’s 16 regions. The two ISP records identified in 2018 and 2019 suggest that co-producing isolates were present before this profile became recurrent in outbreak surveillance, although differences between the two surveillance systems preclude direct temporal comparisons. Quesille-Villalobos et al. (2025) (Quesille-Villalobos et al., 2025) also identified KPC/NDM co-producers causing invasive infections in multiple hospitals and across four species, providing independent evidence of multispecies dissemination.

The concurrent expansion in isolate numbers, host range, carbapenemase profiles, and geographic representation motivated genome-and plasmid-level analyses of the mechanisms underlying this diversification.

### Hybrid sequencing resolved the chromosomes and plasmid repertoires of 30 selected isolates

To move from national surveillance patterns to genome-resolved characterization, we selected 30 isolates for hybrid sequencing to cover different years, bacterial taxa, geographic regions, and carbapenemase profiles. The selected isolates spanned 2018–2024 and represented 11 species from six genera, the three predominant carbapenemase-family combinations, and 19 assigned species-specific sequence type profiles, including recurrent and singleton lineages. This selection strategy supported phylogenomic, resistome, and mobilome analyses across a broad range of biological backgrounds, prioritizing breadth over reproduction of the relative frequencies observed among the 414 surveillance records.

Hybrid sequencing generated consistently high-quality assemblies for all 30 isolates (Table S3). Estimated completeness was 100% for every genome, while contamination ranged from 0.01% to 2.33% (median, 0.38%). Sequencing depth ranged from 78× to 1,028× (median, 164×). Total assembly sizes ranged from 5.05 to 6.59 Mb (median, 5.47 Mb), GC content from 51% to 60%, and the number of predicted coding sequences from 4,701 to 6,159. Each assembly contained a single circular chromosomal contig of 4.66–6.11 Mb and two to nine circular extrachromosomal replicons, yielding 149 complete plasmids across the collection.

Hybrid assembly resolved the genomic context of co-occurring carbapenemases, which is often obscured in short-read assemblies by repetitive regions and homologous plasmid sequences (Arredondo-alonso et al., 2017; Bouras et al., 2024). Complete replicons allowed resistance genes to be assigned to chromosomes or individual plasmids and enabled structural comparison of plasmid backbones, integrons, and repeated resistance modules. This resolution distinguished co-localization of two carbapenemases on a single plasmid from their carriage on separate replicons, two epidemiologically distinct configurations. The closed genomes also connected clonal relationships with the distribution of plasmids and resistance modules across bacterial hosts.

### Taxonomic and phylogenomic breadth supports multiple routes to carbapenemase co-production

The 30 sequenced isolates represented 11 species from six genera (Fig. 2; Figs. S1–S11). *K. pneumoniae* and *Enterobacter hormaechei* were the most frequent species, with eight isolates each, followed by *Citrobacter freundii* with four. Multilocus sequence typing assigned 26 isolates to 19 species-specific sequence type (ST) profiles, while four remained unassigned. Five profiles recurred: *K. pneumoniae* ST11 (n = 3), *E. hormaechei* ST114 (n = 3), *E. hormaechei* ST45 (n = 2), *C. freundii* ST62 (n = 2), and *Klebsiella aerogenes* ST93 (n = 2). The other 14 typed isolates represented singleton profiles, indicating that recurrence was limited to a few bacterial backgrounds.

**Figure 2.**
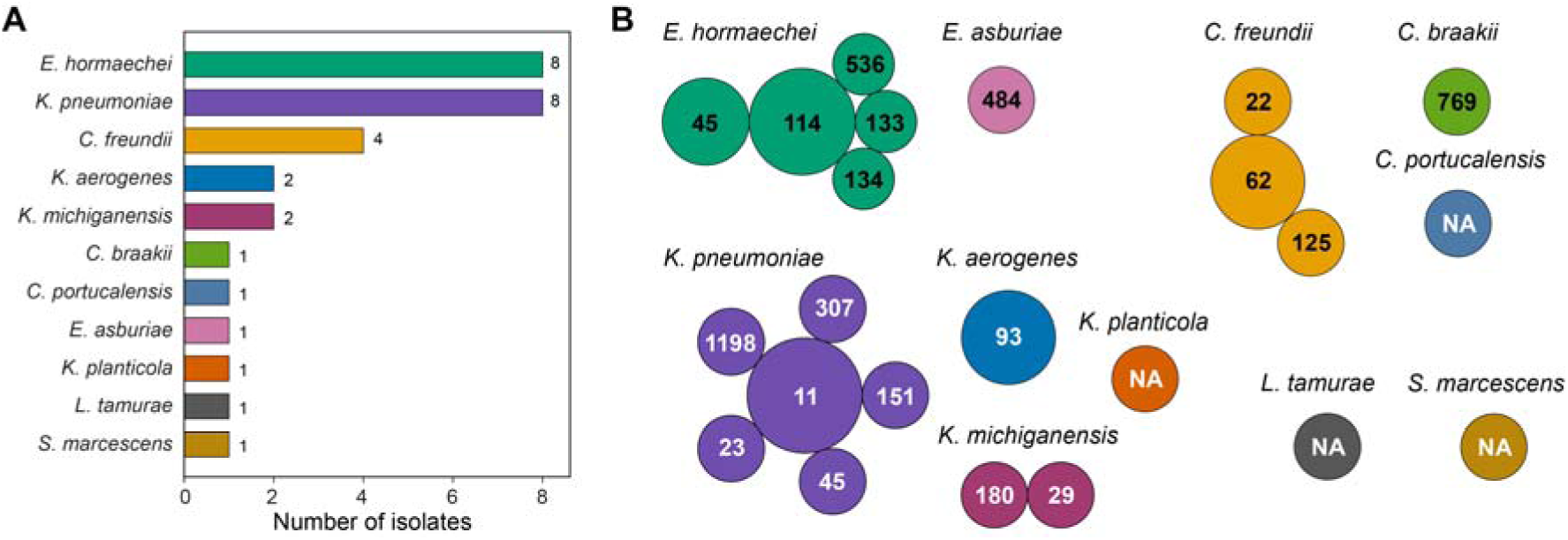
Taxonomic and clonal diversity of the sequenced collection of multiple-carbapenemase-producing Enterobacterales. Thirty isolates collected in Chile between 2018 and 2024 were classified using genome-based taxonomic assignment and species-specific multilocus sequence typing. (A) Taxonomic composition of the collection, showing the number of isolates assigned to each species. (B) Distribution of isolates among species-specific sequence types (STs). Each circle represents one ST within the indicated species; circle area reflects the number of isolates assigned to that ST, and the number within the circle indicates the ST designation. NA denotes isolates without an assigned ST. Colors indicate bacterial species. Because MLST schemes and numerical ST designations are species-specific, ST recurrence was evaluated independently within each species.

The eight *K. pneumoniae* isolates belonged to six STs: ST11, ST23, ST45, ST151, ST307, and ST1198. Among 120 public genomes representing these STs, 61 carried at least one carbapenemase, but only one carried multiple carbapenemases, compared with all eight Chilean isolates (Fig. S1). ST11 and ST307 are established epidemic lineages associated with carbapenemase dissemination in Europe and the Americas, whereas ST23 is a well-recognized hypervirulent lineage (Peirano et al., 2020; Saavedra et al., 2021; van der Zwaluw et al., 2020). The public ST11 comparison was dominated by Colombian genomes (37/39), of which 36 carried KPC-3. In contrast, the three Chilean ST11 isolates carried KPC-66/NDM-1, KPC-2/NDM-1, and KPC-14/NDM-5. Public ST151 genomes originated from Norway, the United States, Singapore, Spain, and Colombia. Eight of 18 carried either KPC-2 or OXA-48, whereas the Chilean ST151 isolate carried KPC-3/NDM-5. These contrasts show that the same clonal backgrounds can acquire distinct carbapenemase complements. ST25 was absent from the sequenced subset despite the persistence of the high-risk ST25-K2/CG10224 lineage in Chile (Veloso et al., 2023). Because ST information was unavailable during isolate selection, the subset was not intended to reproduce the relative frequencies of specific *K. pneumoniae* lineages in national surveillance.

The eight *E. hormaechei* isolates belonged to ST114, ST45, ST133, ST134, and ST536. Among 383 public genomes, 254 carried at least one carbapenemase and 22 carried multiple carbapenemases (Fig. S2). The three Chilean ST114 isolates carried VIM-1/NDM-7, whereas the two ST45 isolates combined VIM-1 with either NDM-7 or KPC-2. ST114 is a globally distributed multidrug-resistant nosocomial lineage involved in the dissemination of carbapenemase-bearing plasmids (Macesic et al., 2023). Fourteen of 16 public ST114 genomes from Switzerland, the United States, Australia, and China carried a carbapenemase, most frequently OXA-48, a profile distinct from VIM-1/NDM-7 in the Chilean isolates. ST133 was also strongly associated with carbapenemases: 30 of 33 public genomes were positive, predominantly for IMP-1, and most originated from Japan. A recent Chilean study identified a closely related ST133 cluster carrying NDM-7 and VIM-1 on separate plasmids (Solar et al., 2026), whereas the ST133 isolate examined here carried NDM-7/KPC-2. ST45, ST134, and ST536 were less extensively represented in public collections but also occurred in carbapenemase-positive backgrounds. Recurrence of ST114 and ST45, combined with variation in their carbapenemase profiles, supports contributions from both clonal persistence and acquisition of different resistance elements.

The *Citrobacter* subset comprised four *C. freundii* isolates assigned to ST22, ST62, and ST125, one *C. braakii* ST769 isolate, and one unassigned *C. portucalensis* isolate. The two *C. freundii* ST62 isolates and the ST125 isolate carried KPC-2/NDM-7, whereas the ST22 isolate carried NDM-7/VIM-2. Among 508 public *C. freundii* genomes, 386 carried at least one carbapenemase and 71 carried multiple carbapenemases (Fig. S3). ST22 was widely represented and strongly associated with carbapenemases, whereas ST62 and ST125 were less frequent. ST22 has been recovered from patients, hospital environments, and wastewater-associated reservoirs, including isolates carrying multiple carbapenemases (Castilla et al., 2005; Hamerlinck et al., 2023; Ju et al., 2025). Quesille-Villalobos et al. (Quesille-Villalobos et al., 2025) also identified *C. freundii* among Chilean KPC/NDM co-producers. The occurrence of NDM-7/VIM-2 in ST22 and KPC-2/NDM-7 across ST62 and ST125 shows that co-production in *C. freundii* spans multiple lineages and enzyme combinations.

The *C. braakii* ST769 isolate carried KPC-2/NDM-7. Of 212 public *C. braakii* genomes, 65 carried a carbapenemase and only six carried multiple carbapenemases (Fig. S4). The four public ST769 genomes originated from Mexico, the United States, and China, and the two carbapenemase-positive representatives carried single GES-5 or IMP-4 enzymes. The *C. portucalensis* isolate VA3495 carried VIM-2/KPC-2. Among 201 public genomes, 108 carried at least one carbapenemase and only two carried multiple carbapenemases (Fig. S5). Neither species was represented in the Chilean KPC/NDM collection described by Quesille-Villalobos et al. (2025). In Latin America, *C. portucalensis* has been recovered from human, animal, and environmental sources, including NDM-1-producing clinical isolates from Brazil (C. H. Camargo et al., 2024; Sellera et al., 2022). These Chilean isolates expand the species and carbapenemase combinations documented within the genus.

The *Enterobacter asburiae* ST484 isolate carried VIM-2/NDM-7. Among 328 public genomes, 118 carried at least one carbapenemase but only three carried multiple carbapenemases (Fig. S6). Fifteen of 26 public ST484 genomes carried a carbapenemase, most frequently IMP-1, KPC-2, or NDM-1, but none reproduced the VIM-2/NDM-7 profile. VIM-1-producing ST484 isolates have also been recovered from hospital and municipal wastewater in the Czech Republic (Sukkar et al., 2024).

Both *K. aerogenes* ST93 isolates carried KPC-2/NDM-7. ST93 was the most frequent lineage in the public collection and included genomes from North America, Brazil, Europe, and Asia, consistent with its international distribution (Morgado et al., 2024). Nevertheless, only 13 of 134 public *K. aerogenes* genomes carried a carbapenemase, none carried multiple carbapenemases, and only one of 34 public ST93 genomes was carbapenemase-positive (Fig. S7). KPC-2/NDM-7 co-production in two Chilean ST93 isolates therefore represents an unusual resistance configuration within this widespread lineage.

The two *K. michiganensis* isolates carried KPC-2/NDM-7 but belonged to ST29 and ST180. Carbapenemases occurred in 183 of 424 public genomes, although only ten carried multiple carbapenemases (Fig. S8). ST29 was represented mainly by Japanese genomes and was frequently associated with IMP-6. ST180 included genomes from Ecuador, North America, Europe, and Asia carrying single carbapenemases such as NDM-1, VIM-1, or KPC-2. Neither lineage reproduced the KPC-2/NDM-7 profile of the Chilean isolates, further showing that members of the *K. oxytoca* species complex can acquire different carbapenemase combinations across multiple bacterial backgrounds.

The *Raoultella planticola*, *Leclercia tamurae*, and *Serratia marcescens* isolates were recovered from human infections and carried KPC-2/NDM-7. Carbapenemases were detected in 53 of 124 public *R. planticola* genomes, 42 of 122 public *Leclercia* genomes, and 83 of 403 public *S. marcescens* genomes, whereas multiple carbapenemases occurred in only five, zero, and one genome, respectively (Figs. S9–S11). Because few *L. tamurae* genomes were available, its phylogenomic context was evaluated against the broader genus *Leclercia*. Carbapenemase-positive *Leclercia* genomes frequently originated from Mexico and were dominated by NDM-1, but none carried multiple carbapenemases. In Chile, KPC-2-producing *S. marcescens* had previously been described in a hospitalized patient, and national outbreak surveillance subsequently documented KPC/NDM co-production in this species (Lara et al., 2026; Quezada-Aguiluz et al., 2020). The completely resolved *S. marcescens* isolate examined here provides genomic and plasmid-level context for this emerging national phenotype.

These isolates also illustrate the extension of clinically important resistance into opportunistic hosts that receive less attention in genomic surveillance. *Raoultella* species occur widely in water, soil, and plants but are increasingly recognized as human pathogens (Sękowska, 2017). *Leclercia* is similarly associated with soil, water, food, and plant environments, and *L. tamurae* was only recently described from isolates recovered from the oak rhizosphere (Maddock et al., 2022). Although human infections caused by *Leclercia* remain uncommon, isolates carrying multidrug-resistance determinants and conjugative plasmids have increasingly been reported (Yescas-Zazueta et al., 2024; Zayet et al., 2021). The *L. tamurae* isolate examined here is therefore notable because it originated from a human infection and carried KPC-2 and NDM-7 on plasmid lineages also circulating among established clinical pathogens in this collection. This finding illustrates how cross-species plasmid dissemination can introduce multidrug resistance into uncommon opportunistic hosts.

The environmental associations of these taxa may facilitate their introduction into water-associated healthcare reservoirs. Hospital water systems, drains, and biofilms can harbor antimicrobial-resistant Gram-negative bacteria and contribute to healthcare-associated transmission (Hayward et al., 2020). This risk is particularly well documented for *S. marcescens*, which can persist on hospital surfaces and in moist environments and has caused outbreaks involving strains with reduced susceptibility to disinfectants (Hanczvikkel et al., 2024). The occurrence of biocide-and metal-associated resistance determinants in the present collection raises the possibility that non-antibiotic selective pressures contribute to the maintenance of these organisms and their plasmids in healthcare environments (Pal et al., 2015). However, environmental persistence and reduced disinfectant susceptibility were not evaluated phenotypically and cannot be inferred from gene carriage alone. These observations identify uncommon environmental and opportunistic Enterobacterales as potential reservoirs of mobile resistance determinants and sources of difficult-to-control nosocomial infections.

Overall, the coexistence of globally disseminated clones, recurrent local lineages, and uncommon opportunistic hosts argues against emergence through a single clonal event. Instead, the phylogenomic patterns support both clonal persistence and repeated acquisition of carbapenemase determinants across multiple Enterobacterales taxa. We therefore examined the plasmidome to determine whether related resistance platforms circulated across these bacterial backgrounds.

### Broad β-lactam resistance is accompanied by diverse accessory resistomes and potential co-selection signatures

All 30 isolates were resistant to cefotaxime, meropenem, and imipenem (Table S2). Cefepime results were interpretable for 28 isolates, ceftazidime for 23, and ertapenem for 29; every isolate with an interpretable result was resistant. Tigecycline measurements were available for 28 isolates, but no interpretive categories were applied. Overall, the collection displayed a uniform broad-spectrum β-lactam resistance phenotype consistent with the carriage of multiple carbapenemases.

The 30 genomes contained 82 distinct antimicrobial resistance determinants, with individual isolates carrying between 5 and 26 (median, 15). Resistance-gene burden showed both taxonomic and lineage-associated structure. *K. pneumoniae* had the highest median number of determinants (20.5). The three ST11 isolates carried between 20 and 26 determinants but differed in their accessory gene complements, indicating substantial within-lineage variation. The clearest within-species contrast occurred in *C. freundii*: the ST22 and ST125 isolates carried 23 and 24 determinants, respectively, whereas the two ST62 isolates shared the same 11-gene profile. The three *E. hormaechei* ST114 isolates had similar burdens of 14 to 16 determinants. In contrast, the two *K. aerogenes* ST93 isolates carried 13 and 19 determinants despite sharing the same KPC-2/NDM-7 combination. Resistome variation therefore reflected both lineage-associated repertoires and differences in accessory resistance content (Fig. 3A,C).

**Figure 3.**
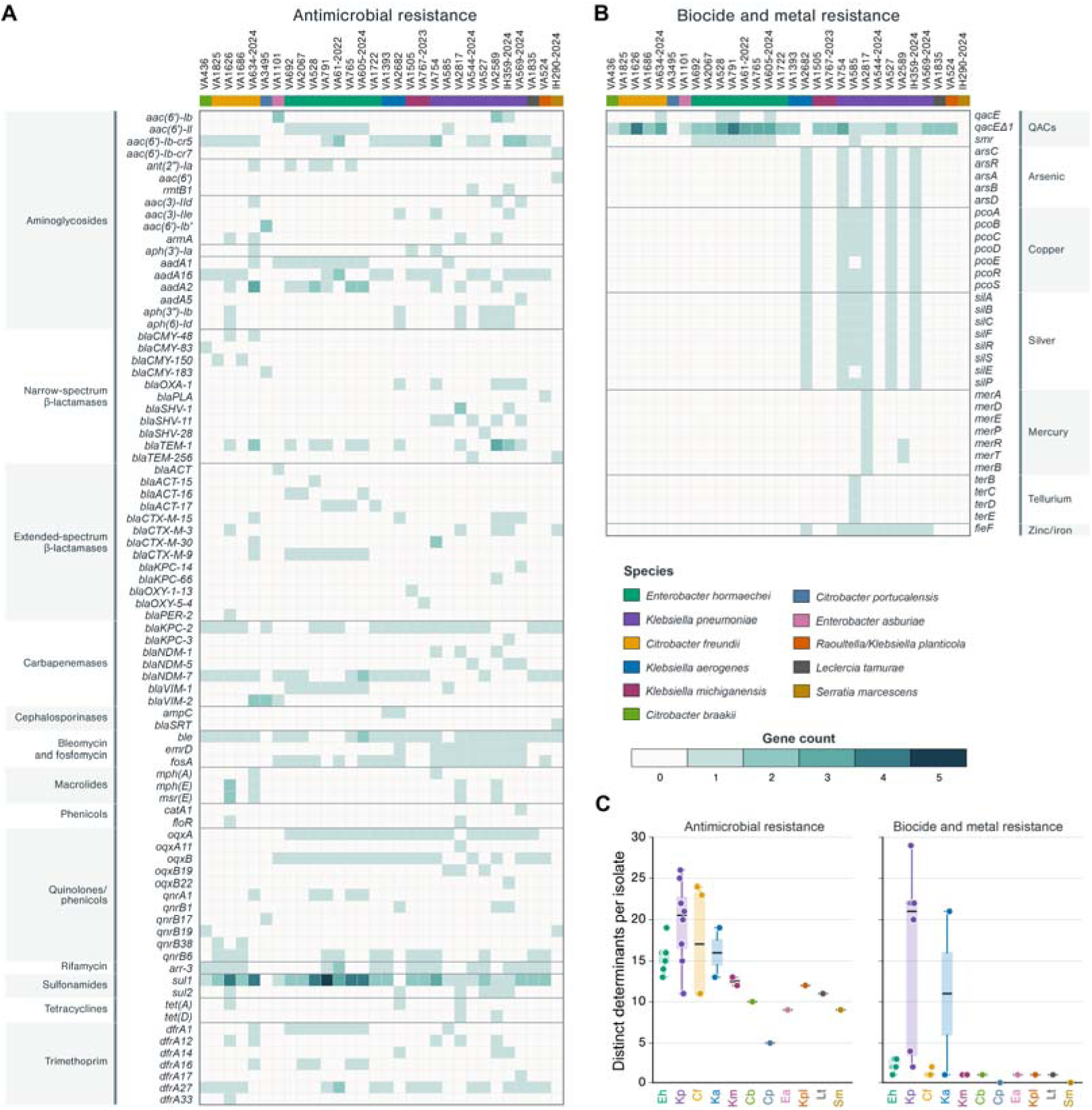
Distribution and species-level variation in antimicrobial, biocide, and metal resistance determinants across the 30 sequenced isolates. (A) Heatmap showing the presence and copy number of 82 antimicrobial resistance determinants, grouped according to antimicrobial class. (B) Heatmap showing 35 determinants associated with resistance or tolerance to quaternary ammonium compounds, arsenic, copper, silver, mercury, tellurium, and zinc/iron. In (A) and (B), columns represent isolates and rows represent individual genes or alleles. The upper annotation strip indicates bacterial species, and cell color represents the number of copies detected per genome (0–5). (C) Species-stratified distributions of the number of distinct antimicrobial resistance determinants (left) and biocide-or metal-associated determinants (right) detected per isolate. Points represent individual isolates; boxes indicate the interquartile range, central lines indicate medians, and whiskers extend to the most extreme values within 1.5 times the interquartile range. Species represented by only one or two isolates are displayed primarily by their individual data points. Species abbreviations: Eh, *Enterobacter hormaechei*; Kp, *Klebsiella pneumoniae*; Cf, *Citrobacter freundii*; Ka, *Klebsiella aerogenes*; Km, *Klebsiella michiganensis*; Cb, *Citrobacter braakii*; Cp, *Citrobacter portucalensis*; Ea, *Enterobacter asburiae*; Kpl, *Raoultella/Klebsiella planticola*; Lt, *Leclercia tamurae*; and Sm, *Serratia marcescens*. QACs, quaternary ammonium compounds.

β-Lactam and aminoglycoside resistance genes were present in every isolate, whereas sulfonamide and trimethoprim resistance determinants occurred in 28 and 27 isolates, respectively. Twenty-three isolates carried *qnr* alleles, and 20 carried *aac(6*′*)-Ib-cr*, which can contribute to reduced susceptibility to both aminoglycosides and fluoroquinolones. These determinants were distributed across multiple genera, although individual alleles sometimes showed narrower associations. For example, *qnrB38* was restricted to the two *C. freundii* ST62 isolates. In contrast, *fosA* showed a marked taxonomic distribution: it occurred in all eight *K. pneumoniae* isolates, both *K. aerogenes* isolates, and six of eight *E. hormaechei* isolates, but was absent from all *Citrobacter* genomes. Other frequent determinants included *arr-3* (n = 17), while genes associated with macrolide, tetracycline, and phenicol resistance were less common. Because susceptibility to these antimicrobial classes was not evaluated consistently, these findings indicate genomic resistance potential rather than confirmed phenotypic resistance.

The β-lactam resistome comprised nine carbapenemase alleles spanning three families: KPC (KPC-2,-3,-14, and-66), NDM (NDM-1,-5, and-7), and VIM (VIM-1 and-2). Each isolate carried two distinct alleles, yielding ten combinations. KPC/NDM combinations predominated (n = 19), followed by NDM/VIM (n = 6) and KPC/VIM (n = 5). KPC-2/NDM-7 was the most frequent combination (n = 13), whereas KPC-2/VIM-1 and NDM-7/VIM-1 occurred in four isolates each. Additional ESBL, AmpC-type, and broad-spectrum β-lactamases included CTX-M, PER-2, CMY, ACT, OXA, SHV, and TEM variants. This combined repertoire was consistent with the uniform β-lactam resistance phenotype, although the contribution of individual enzymes could not be resolved in these complex multidrug-resistant backgrounds.

Three isolates carried an additional copy of one carbapenemase allele, bringing the total to 63 gene copies. Of these, 62 (98.4%) were plasmid-borne; the only chromosomal copy was *bla*_NDM-1_ in *K. pneumoniae* VA754. Plasmids therefore accounted for nearly the entire carbapenemase repertoire. The complete replicons distinguished the additional copies as intraplasmid duplications or redundant carriage on co-resident plasmids, as examined below.

Colistin displayed a less direct genotype-phenotype relationship. Interpretable results were available for 28 isolates: 26 were classified as intermediate and two as resistant. No result was available for *E. hormaechei* VA791, while interpretation was not applicable to *S. marcescens* IH290 because this species is intrinsically resistant to colistin. The two resistant isolates, *K. aerogenes* VA1393 and *K. pneumoniae* VA2817, had MICs >4 µg/mL. Three isolates carried plasmid-encoded *mcr-9.2*. Of the two with susceptibility results, *E. hormaechei* VA528 and VA765 had MICs ≤1 µg/mL and were classified as intermediate; VA791 was the third *mcr-9.2*-positive isolate. Neither colistin-resistant isolate carried *mcr-9.2*. Thus, *mcr-9.2* carriage did not correspond to phenotypic resistance in the available data, while the mechanisms responsible for resistance in VA1393 and VA2817 remain unresolved.

Gene co-occurrence revealed several conserved resistance modules. All 25 NDM-positive isolates carried *ble*, which was absent from the five NDM-negative isolates, supporting conservation of an NDM-*ble* module across species and NDM alleles. Likewise, *sul1* and *qacE*Δ*1* co-occurred in 27 isolates, reflecting their characteristic association within the conserved 3′ region of class 1 integrons (Partridge et al., 2018). Other recurrent combinations included *arr-3*/*dfrA27* in 17 isolates from multiple genera and *armA*/*mph(E)*/*msr(E)* in four isolates with comparatively large resistomes, comprising two *K. pneumoniae* and two *C. freundii* from different ST backgrounds. Copy-number variation was also evident, with up to five copies of *sul1* and four of *qacE*Δ*1* in individual isolates. Complete assemblies verified that these copies formed fully resolved repeated regions and enabled their genomic contexts to be examined.

Determinants associated with reduced susceptibility to quaternary ammonium compounds were particularly widespread, with at least one of *qacE*, *qacE*Δ*1*, or *smr* detected in 27 isolates. Their linkage to antimicrobial resistance genes within integrons and other mobile regions provides a plausible mechanism for co-selection, whereby exposure to either antibiotics or disinfectants may favor maintenance of the entire region (Boyce, 2023; Murray et al., 2024; Pal et al., 2015). This association may be especially relevant in healthcare environments, where sublethal disinfectant exposure and contaminated aqueous reservoirs have been linked to the persistence and transmission of multidrug-resistant Gram-negative bacteria, including nosocomial outbreaks caused by *S. marcescens* (Boyce, 2023; Hanczvikkel et al., 2024).

Nine isolates also carried at least one selected metal-associated resistance system. Complete or near-complete *pco* and *sil* loci occurred in six isolates, *ars* genes in five, *mer* genes in two, and a *ter* cluster in one (Fig. 3B,C). Their co-occurrence with antimicrobial resistance genes provides an additional genomic basis for potential co-selection by metals or other non-antibiotic stressors.

Overall, the isolates shared extensive β-lactam resistance but differed markedly in their accessory resistance repertoires. The frequent association of antibiotic-, biocide-, and metal-resistance determinants suggests that some mobile resistance regions may be maintained under multiple selective pressures, although their phenotypic relevance remains to be established. The complete assemblies subsequently allowed the genomic linkage, structural organization, and mobility of these determinants to be examined.

### A diverse plasmidome with high predicted transfer potential is structured around three recurrent lineages

The 149 complete plasmids recovered from the 30 isolates corresponded to two to nine plasmids per genome (median, four), with a median size of 45.1 kb and a maximum of 375.6 kb (Table S4). MOB-typer classified 72 plasmids as conjugative (48.3%), 33 as mobilizable (22.1%), and 44 as non-mobilizable (29.5%). Median size differed markedly among these classes, at 58.4, 5.2, and 4.4 kb, respectively (Fig. 4A). Carbapenemase-positive plasmids had a higher median size than carbapenemase-negative plasmids (55.0 versus 6.9 kb), and 55 of 57 (96.5%) were conjugative (n = 49) or mobilizable (n = 6). Carbapenemase carriage was therefore concentrated in larger plasmids with predicted transfer potential, supporting their role as plausible vehicles for horizontal dissemination, although experimental confirmation is required.

**Figure 4.**
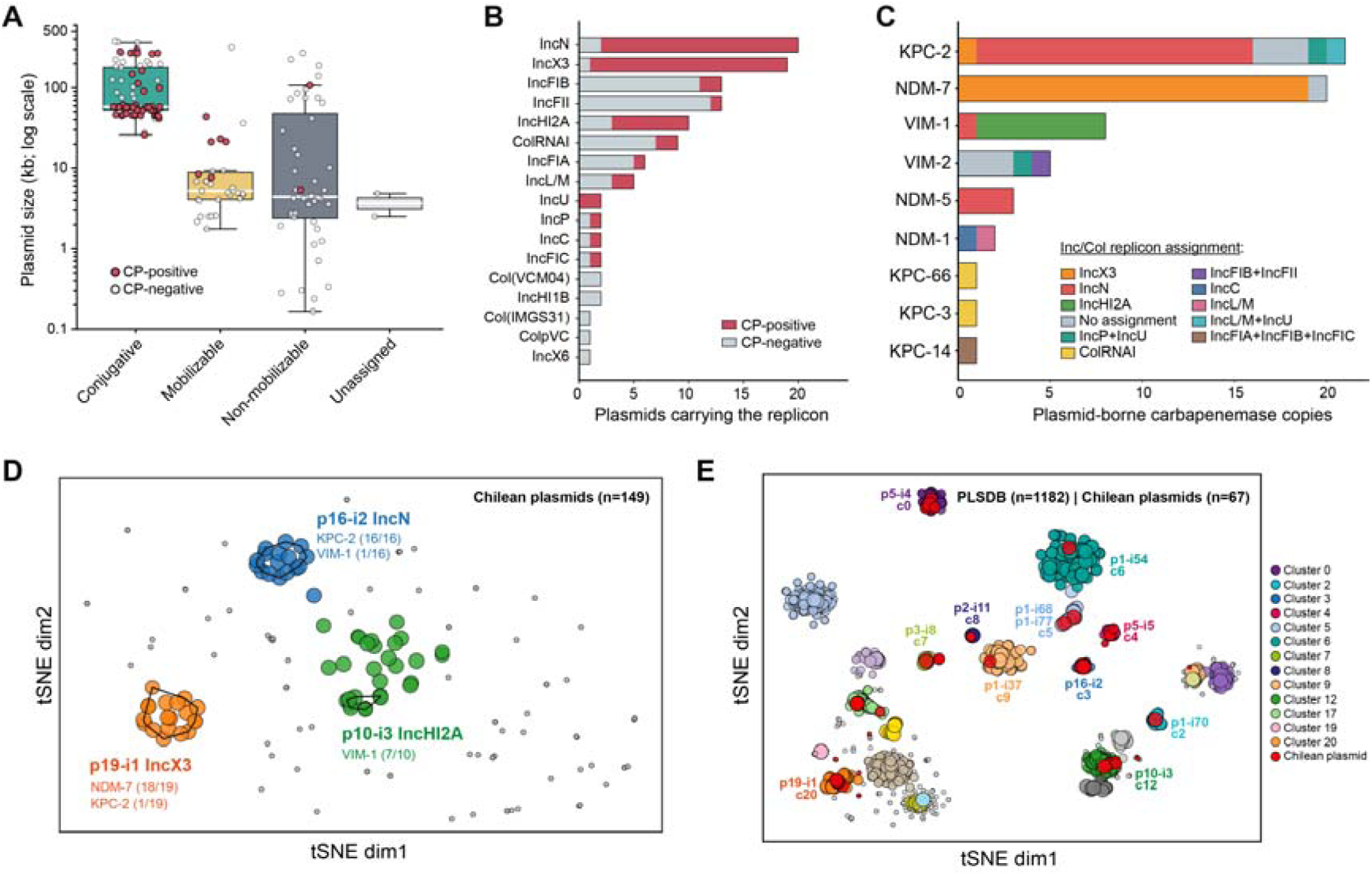
Mobility, replicon composition, carbapenemase content, and sequence-based organization of the complete plasmidome. (A) Plasmid size distributions according to MOB-typer mobility class. Filled red and open circles represent carbapenemase-positive and carbapenemase-negative plasmids, respectively. Boxes indicate the median and interquartile range; whiskers extend to the most extreme values within 1.5 times the interquartile range. (B) Conventional Inc/Col replicon assignments stratified by carbapenemase content. Replicon categories are non-exclusive, and multireplicon plasmids contribute to each detected category. (C) Distribution of plasmid-borne carbapenemase copies according to the complete Inc/Col replicon profile of the corresponding plasmid. Each carbapenemase copy is counted once, and multireplicon profiles are retained. “No assignment” includes plasmids with only MOB-suite rep_cluster assignments or no detected replicon. (D) MGEcluster t-SNE projection of the 149 complete plasmids reconstructed from the Chilean isolates. Colors indicate HDBSCAN-defined groups; outlines and labels identify the three predominant study-specific plasmid clusters. (E) MGEcluster t-SNE projection of 67 Chilean plasmids assigned to MOB-cluster pairs represented among the 57 carbapenemase-positive plasmids, together with 1,182 complete PLSDB plasmids carrying the same primary and secondary MOB-cluster pairs. The Chilean set therefore included the 57 carbapenemase-positive plasmids and ten carbapenemase-negative members of the same MOB-defined groups. Selected Chilean plasmid clusters are labelled, with the three predominant lineages highlighted.

Replicon typing revealed substantial diversity (Fig. 4B). IncN and IncX3 were the most frequent incompatibility groups, occurring in 20 and 19 plasmids, respectively, followed by IncFIB and IncFII (n = 13 each), IncHI2A (n = 10), ColRNAI (n = 9), IncFIA (n = 6), and IncL/M (n = 5). Assignments were non-exclusive because several plasmids carried multiple replicon types. Carbapenemase-positive plasmids accounted for 18 of the 20 IncN plasmids, 18 of the 19 IncX3 plasmids, and seven of the ten IncHI2A plasmids, whereas most IncFIB and IncFII plasmids lacked carbapenemases. Conventional Inc/Col typing did not resolve the entire collection, as 32 plasmids lacked a detectable replicon assignment and another 25 were classified only through MOB-suite rep_cluster identifiers.

The 62 plasmid-borne carbapenemase copies showed marked but non-exclusive associations with particular replicon backgrounds (Fig. 4C). Nineteen of the 20 *bla*_NDM-7_ copies occurred on IncX3 plasmids, 15 of the 21 *bla*_KPC-2_ copies on IncN plasmids, and seven of the eight *bla*_VIM-1_ copies on IncHI2A plasmids. All three *bla*_NDM-5_ copies were also associated with IncN. The remaining copies occurred in less frequent or unassigned profiles, including IncP/IncU, IncFIB/IncFII, IncC, IncL/M, ColRNAI, and multireplicon IncF plasmids. The predominant alleles were therefore concentrated in a small number of replicon backgrounds but were not restricted to them.

For greater resolution, plasmids with identical primary and secondary MOB-cluster assignments were classified into groups designated pA-iB, where A indicates group size and B its sequential identifier after ordering groups from largest to smallest. Of the 149 plasmids, 147 were assigned to 77 groups and two remained untyped. Most groups were sparsely represented: 55 were singletons, 14 contained two plasmids, three contained three, and two contained five.

Three groups were markedly more frequent than the remainder. The IncX3 p19-i1, IncN p16-i2, and IncHI2A-associated p10-i3 lineages comprised 19, 16, and 10 plasmids, respectively, together accounting for 45 of the 149 plasmids (30.2%). All were classified as conjugative. Carbapenemases occurred in 18 of 19 p19-i1 plasmids, all 16 p16-i2 plasmids, and seven of ten p10-i3 plasmids, with predominant associations with *bla*_NDM-7_, *bla*_KPC-2_, and *bla*_VIM-1_, respectively. The remaining plasmidome consisted largely of low-frequency groups and singletons, including several alternative carbapenemase vehicles.

Available MOB-typer outputs were consistent within each major lineage. MOB-typer integrates host distributions from its reference plasmid database with curated literature evidence to estimate an overall replicative host range (Robertson et al., 2020). This prediction converged at the order Enterobacterales for p19-i1 and p10-i3 and at the broader class Gammaproteobacteria for p16-i2. The NCBI-observed ranges matched these overall predictions for every member of the three lineages. Literature-derived assignments converged at the family Enterobacteriaceae for all p19-i1 members and at the order Enterobacterales for all p10-i3 members and 15 of the 16 p16-i2 members.

Because MOB-cluster assignments are reference-dependent and public plasmid collections remain geographically uneven, with limited South American representation compared with China, North America, and Europe (A. P. Camargo et al., 2024; Schmartz et al., 2022), we evaluated the coherence of these groups using whole-plasmid sequences. MGEcluster analysis followed by t-SNE projection and HDBSCAN identified three density-defined groups containing 19, 17, and 30 plasmids, while 83 remained ungrouped under the selected parameters (Fig. 4D). The 19 p19-i1 members coincided exactly with the 19-member sequence-defined group, and all 16 p16-i2 plasmids occurred within the 17-member group. In contrast, the ten p10-i3 plasmids formed a more narrowly defined lineage within the broader 30-member group.

To determine whether the Chilean plasmid lineages associated with carbapenemase carriage were also represented among publicly available plasmids, we constructed a targeted comparison with PLSDB. The 57 carbapenemase-positive plasmids were used to define the relevant primary-secondary MOB-cluster pairs. We then included all Chilean plasmids assigned to these pairs, regardless of carbapenemase carriage, allowing the comparison to capture related backbones with different resistance cargo. This yielded 67 Chilean plasmids, including ten carbapenemase-negative members of the same MOB-defined groups. Exact-pair matching recovered 1,182 complete PLSDB plasmids, producing a combined dataset of 1,249 sequences for whole-plasmid analysis (Fig. 4E).

The major Chilean lineages differed in their relationships with public sequences. All 16 p16-i2 plasmids formed a sequence-defined group with ten PLSDB plasmids, whereas eight of the ten p10-i3 plasmids grouped with 64 public sequences and two remained ungrouped. In contrast, the 19 p19-i1 plasmids were distributed among three HDBSCAN groups containing public plasmids, while seven remained ungrouped. Thus, p16-i2 and p10-i3 were closely associated with plasmid populations represented in PLSDB, whereas the shared AA038/AH615 assignment of p19-i1 encompassed greater whole-sequence heterogeneity.

The predominance of plasmid groups represented in only one or two isolates suggested that the 30-isolate panel captured only part of the underlying plasmid repertoire. We therefore used lineage-accumulation and richness analyses to evaluate sampling completeness and estimate the potential for discovering additional lineages. For this analysis, plasmids sharing the same primary-secondary MOB-cluster pair were treated as members of the same operational lineage. Among the 147 typed plasmids, 78 lineages were observed. Fifty-eight occurred in a single isolate and 12 in two isolates, such that 70 of 78 lineages (89.7%) were detected in no more than two bacterial hosts (Fig. S12). The accumulation curve remained far from a plateau after 30 isolates. The bias-corrected Chao2 estimator yielded an asymptotic richness estimate of approximately 201 operational lineages, with an estimated incidence coverage of 60.8%. These results support substantial potential for additional lineage discovery within the complete plasmid repertoire of multiple-carbapenemase-producing Enterobacterales.

Broader local and regional studies based on complete, plasmid-resolved genomes will be required to characterize this diversity. Such studies remain challenging in South America and other low-and middle-income settings because of sequencing costs, infrastructure, reagent supply chains, and limited access to specialized expertise (NIHR Global Health Research Unit on Genomic Surveillance of AMR, 2020; Calero-Cáceres et al., 2023). These constraints are especially consequential for plasmid surveillance, which generally requires long-read or hybrid sequencing to reconstruct complete replicons (Bouras et al., 2024) and contribute to the underrepresentation of South American plasmids in public databases (Schmartz et al., 2022).

Together, these analyses identified three recurrent conjugative lineages that concentrated the most frequent carbapenemase alleles within a highly diverse and incompletely sampled plasmidome. Their recurrence supports dissemination of successful plasmid backbones, whereas the numerous low-frequency groups and singletons indicate that carbapenemase co-production also involves a broad reservoir of less prevalent architectures. We therefore examined the host distribution, conserved backbone, and accessory gene repertoire of the major lineages and selected lower-frequency groups.

### Major plasmid lineages combine conserved backbones with modular accessory variation

Pan-plasmidome analysis using Panaroo revealed conserved backbones accompanied by variably distributed accessory genes in p19-i1, p16-i2, and p10-i3 (Fig. S13). Representative architectures are shown in Fig. 5A–C. The lineages differed in host distribution, backbone conservation, size, and resistance-gene content.

**Figure 5.**
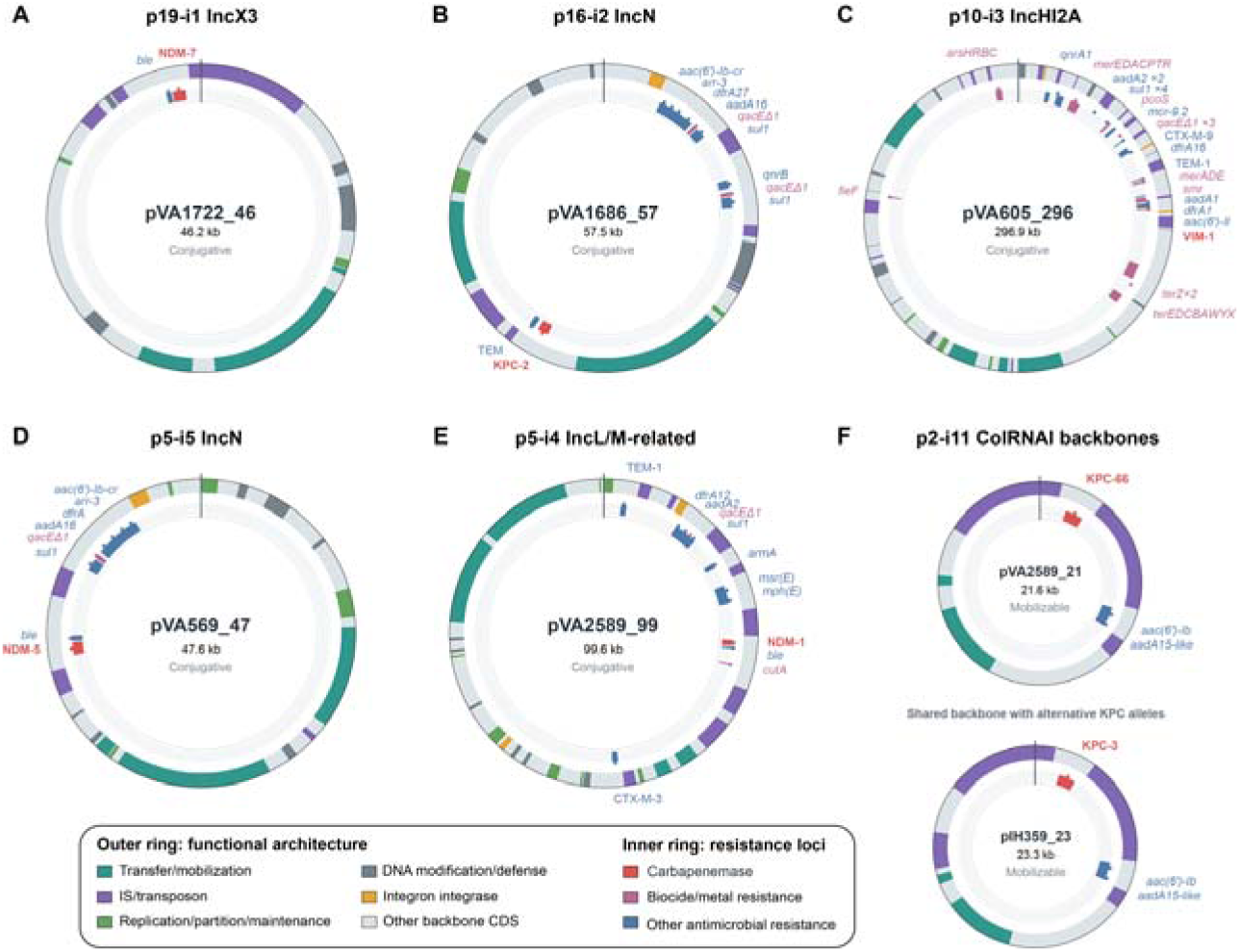
Representative backbone architectures of recurrent and selected lower-frequency carbapenemase plasmid lineages. Circular maps show representatives of the three recurrent plasmid lineages: (A) p19-i1 IncX3, pVA1722_46; (B) p16-i2 IncN, pVA1686_57; and (C) p10-i3 IncHI2A, pVA605_296. Lower-frequency lineages are represented by (D) p5-i5 IncN, pVA569_47; (E) p5-i4 IncL/M-related, pVA2589_99; and (F) two p2-i11 ColRNAI plasmids with a shared backbone carrying alternative KPC alleles, pVA2589_21 (*bla*_KPC-66_) and pIH359_23 (*bla*_KPC-3_). The outer track classifies coding sequences by functional category, whereas the inner track shows carbapenemases and other antimicrobial-, biocide-, and metal-resistance determinants. Gene labels follow the corresponding resistance-class colors. The radial line at 12 o’clock indicates the sequence origin. Genomic positions are shown relative to each plasmid; circle diameters are not proportional to plasmid size.

The IncX3 p19-i1 lineage had the broadest observed host distribution within the collection, comprising 19 plasmids recovered from six genera and ten species (Table S4; Figs. 5A and S13A). These included *Enterobacter* (n = 6), *Klebsiella* (n = 5), *Citrobacter* (n = 5), and single representatives in *Serratia*, *Raoultella*, and *Leclercia*. The most frequent hosts were *E. hormaechei* (5/19) and *C. freundii* (4/19). Its distribution among multiple species and sequence types supports cross-species circulation of the IncX3 backbone.

The p19-i1 plasmids ranged from 40.9 to 91.0 kb and collectively contained 139 gene clusters, including 46 core (33.1%), 24 shell, and 69 cloud clusters. Eighteen carried *bla*_NDM-7_ together with *ble*. The carbapenemase-negative plasmid pVA1101_40 had a reduced repertoire of 52 clusters, whereas pVA605_91 contained 117 clusters and two copies of *bla*_NDM-7_. pVA2682_52 was the only member carrying *bla*_KPC-2_ in addition to *bla*_NDM-7_. Other accessory differences included arsenic-and mercury-resistance determinants, consistent with the gain or loss of discrete resistance-associated modules within the lineage.

This association between IncX3 plasmids and *bla*_NDM-7_ is consistent with previous Chilean evidence. Quesille-Villalobos et al. (Quesille-Villalobos et al., 2025) identified relatively conserved IncX3/*bla*_NDM-7_ plasmids in dual KPC/NDM-producing isolates from four Enterobacterales species. The present collection extends this association across a broader set of observed bacterial hosts and identifies the corresponding plasmids as a coherent MOB-defined and sequence-defined lineage. However, direct sequence comparison would be required to establish whether the previously reported plasmids belong specifically to p19-i1.

The p16-i2 lineage comprised 16 plasmids recovered from five genera and eight species (Table S4; Figs. 5B and S13B). The most frequent hosts were *E. hormaechei* (4/16), *C. freundii* (3/16), and *K. pneumoniae* (3/16), but the lineage also occurred in *Raoultella* and *Leclercia*. Its hosts included *K. pneumoniae* ST11, ST23, and ST307 and four *E. hormaechei* sequence types. Fifteen plasmids carried an IncN replicon, whereas one lacked a conventional incompatibility assignment; the group is therefore considered IncN-associated rather than uniformly IncN-typed.

p16-i2 had the most conserved gene repertoire of the three lineages. Its members ranged from 41.3 to 61.2 kb and contained 80 gene clusters, including 51 core (63.8%), 23 shell, and six cloud clusters. All carried *bla*_KPC-2_, and four also contained a partial *bla*_TEM_-family sequence that could not be assigned confidently to a specific allele. Most members contained approximately 73 clusters. The reduced plasmids pVA61_41 and pVA528_44 contained 54 and 56 clusters, respectively, while the principal divergent member, pVA585_53, carried an additional integron containing *bla*_VIM-1_, *aadA1*, and *smr*. p16-i2 therefore represents a conserved KPC-2 backbone circulating across diverse Enterobacterales while accommodating occasional deletions and accessory resistance modules.

Evidence for circulation of p16-i2 in Chile extends across studies and sampling periods. An early relative was pKpn-3 (GenBank MT949189.1), an IncN plasmid associated with a multispecies KPC-2 outbreak in a Chilean hospital in 2015, in which *bla*_KPC-2_ was embedded in the non-Tn4401 element NTEKPC-IIe within the transfer region (Wozniak et al., 2021). Comparative analysis identified related platforms in Colombian *K. pneumoniae* and *K. variicola*, together with four Chilean KPC-2 plasmids carrying the AA552/AI756 pair that defines p16-i2 and related to pKpn-3 and the Colombian representatives (Veloso et al., 2023) (Veloso et al., 2023). Additional IncN/KPC-2 plasmids were detected in multiple Chilean species collected during 2019–2022, although their assignment to this exact pair remains undetermined (Quesille-Villalobos et al., 2025). Detection of p16-i2 here in 16 isolates from eight species strengthens evidence for regional persistence and cross-species dissemination, underscoring the value of locally generated plasmid-resolved data. Experimental transfer of pVA585_53 to *E. coli* supports its predicted conjugative capacity (Gálvez-Silva et al., 2024).

The IncHI2A p10-i3 lineage showed a more restricted host distribution (Table S4; Figs. 5C and S13C). Eight of its ten plasmids occurred in *E. hormaechei*, with the remaining two recovered from *L. tamurae* and *C. freundii*. The eight *E. hormaechei* hosts represented five sequence types, excluding expansion of a single clone as the sole explanation for this association. All seven *bla*_VIM-1_-positive members occurred in *E. hormaechei*, whereas the plasmids from *L. tamurae*, *C. freundii*, and the ST133 *E. hormaechei* isolate lacked this carbapenemase. Thus, p10-i3 was preferentially associated with *E. hormaechei*, while *bla*_VIM-1_ was a recurrent but not fixed component of the lineage.

These plasmids were substantially larger than p19-i1 and p16-i2, ranging from 265.9 to 375.6 kb, and contained 523 gene clusters: 202 core (38.6%), 253 shell, and 68 cloud clusters. Their conserved repertoire included replication and transfer functions together with arsenic-and tellurium-resistance systems. The seven *bla*_VIM-1_-positive members shared an accessory module containing *sul1*, *qacE*Δ*1*, *smr*, *aac(6*′*)-Il*, *dfrA1*, and *aadA1*. The two largest plasmids, pVA1722_375 and pVA1835_363, shared 124 clusters absent from the remaining members, including type VI secretion system components, transport and stress-response functions, and numerous insertion sequences and transposases.

The association between *bla*_VIM-1_ and large conjugative IncHI2 plasmids has also been observed outside Chile. IncHI2 plasmids of approximately 270–300 kb have mediated multiclonal and multispecies dissemination of *bla*_VIM-1_ in European hospitals and have been recovered from *E. hormaechei* in non-clinical settings (Coelho et al., 2012; Villa et al., 2014; Sadek et al., 2020). Some carried resistance repertoires resembling those of the *bla*_VIM-1_-positive p10-i3 plasmids, including *bla*_CTX-M-9_, aminoglycoside-resistance genes, *dfrA1*, and *sul1*. These similarities place p10-i3 within a broader family of multidrug-resistance IncHI2 plasmids, although whole-sequence comparisons would be required to establish whether previously reported representatives belong to the same lineage.

The three lineages therefore followed distinct dissemination patterns. p19-i1 combined the broadest observed host distribution with modular accessory variation and a strong association with *bla*_NDM-7_. p16-i2 represented a highly conserved KPC-2 backbone with documented persistence, cross-species dissemination in Chile, and evidence of regional circulation. In contrast, the larger and more variable p10-i3 lineage was preferentially associated with *E. hormaechei* and recurrent *bla*_VIM-1_ carriage.

### Lower-frequency plasmid lineages and exceptional architectures broaden the diversity of carbapenemase carriage

Although p19-i1, p16-i2, and p10-i3 comprised 45 of the 149 reconstructed plasmids, they accounted for 41 of the 57 carbapenemase-positive plasmids (71.9%). The remaining 16 were distributed among lower-frequency groups and singleton backbones (Figs. 5D–F and 6; Table S4). Exceptional members of the predominant lineages also carried dual-carbapenemase configurations or gene duplications. Together, these plasmids introduced alternative backbones, less frequent alleles, and resistance architectures not represented by typical members of the three major lineages.

The p5-i5 group comprised five related plasmids ranging from 17.4 to 51.8 kb (Fig. 5D). Four were recovered from *K. pneumoniae*, whereas the shortest member, pVA791_17, occurred in *E. hormaechei* ST45. Three plasmids carried *bla*_NDM-5_ and were found in *K. pneumoniae* ST151, ST1198, and ST11, indicating dissemination of the resistance-bearing backbone across unrelated clonal backgrounds. The fourth *K. pneumoniae* member carried *bla*_CTX-M-30_, while pVA791_17 lacked an identified β-lactamase. Four members were predicted to be conjugative; pVA791_17 was non-mobilizable, lacked a conventional Inc assignment, and probably represents a reduced derivative. Thus, p5-i5 was not restricted to *K. pneumoniae*, although its association with *bla*_NDM-5_ was confined to this species in the present collection.

p5-i5 was predominantly IncN-associated but differed from p16-i2 in both MOB-cluster assignments. Public plasmids with the same AB161/AJ783 pair have been recovered from other Enterobacterales, including a KPC-2 plasmid from *Citrobacter youngae* in China, showing that this backbone can cross species and acquire alternative carbapenemase modules (Liu et al., 2025). IncN plasmids carrying *bla*_NDM-5_ nevertheless remain comparatively uncommon in *K. pneumoniae*, in which *bla*_NDM-1_ is more frequent and *bla*_NDM-5_ is usually associated with other plasmid backgrounds (Zeng et al., 2022). In contrast, NDM-5 has become the predominant NDM variant in *E. coli* globally, although it remains less frequent than NDM-1 in South American collections (Xia et al., 2024). Reports from Argentina, Brazil, and Uruguay indicate recent emergence of NDM-5-producing *K. pneumoniae* across several high-risk lineages, but do not establish an association with p5-i5 (Alvarez et al., 2024; de Oliveira et al., 2023; Magallanes et al., 2025). The three Chilean p5-i5 plasmids therefore identify a locally relevant vehicle for a still uncommon carbapenemase variant and illustrate how conventional Inc typing can group genetically and epidemiologically distinct plasmid lineages.

The p5-i4 group contained five conjugative plasmids ranging from 84.3 to 112.9 kb and recovered from *K. pneumoniae*, *C. freundii*, and *S. marcescens* (Fig. 5E). pVA2589_99 carried *bla*_NDM-1_ and pVA2817_112 carried *bla*_KPC-2_, while the remaining members encoded different combinations of CTX-M and TEM β-lactamases. Most carried an IncL/M replicon, and one additionally carried IncU. Their AA002/AH532 MOB-cluster pair corresponds to the pCTX-M-3-like IncM2 family, a globally distributed group with conserved transfer backbones and variable multidrug-resistance regions (Kern-Zdanowicz, 2021). A previous Chilean representative, pVA833-92, carried *bla*_KPC-2_ within the novel IS26-flanked NTE_KPC_-IIh platform and was the only AA002/AH532 plasmid identified in the South American collection examined by Veloso et al. (2023). Recovery of p5-i4 from three genera and with NDM-1, KPC-2, or ESBL cargo expands its known local host and resistance-gene diversity and supports its capacity to acquire, replace, or lose β-lactam resistance modules.

The p2-i11 group comprised two mobilizable ColRNAI plasmids of 21.6 and 23.3 kb recovered from *K. pneumoniae* ST11 and ST151 (Fig. 5F). pIH359_23 carried *bla*_KPC-3_, whereas pVA2589_21 carried the less frequent *bla*_KPC-66_ variant. Their shared AA356/AI318 MOB-cluster pair, closely related backbones, and similar carbapenemase environments suggest diversification from a common KPC-bearing ancestor. KPC-66 retains the substitution distinguishing KPC-3 from KPC-2 and contains an additional two-amino-acid deletion within the Ω-loop, making it more plausibly a derivative of KPC-3 than a direct single-step derivative of KPC-2. Consistently, KPC-66 was first identified among ceftazidime-avibactam-resistant variants that evolved from plasmid-borne *bla*_KPC-3_ during antimicrobial treatment (Brunetti et al., 2026; Carattoli et al., 2021). The Chilean configuration is compatible with local allelic diversification after association with the p2-i11 backbone, although independent acquisition cannot be excluded.

Three structurally unrelated plasmids each carried two carbapenemase alleles within a single replicon (Fig. 6A–C). The IncX3 p19-i1 plasmid pVA2682_52 carried *bla*_NDM-7_ and *bla*_KPC-2_, the IncN p16-i2 plasmid pVA585_53 carried *bla*_VIM-1_ and *bla*_KPC-2_, and the IncP/IncU plasmid pVA3495_43 carried *bla*_VIM-2_ and *bla*_KPC-2_. The first two were conjugative and the third mobilizable. These configurations show that two carbapenemases can be carried on the same transferable plasmid rather than on separate replicons.

**Figure 6.**
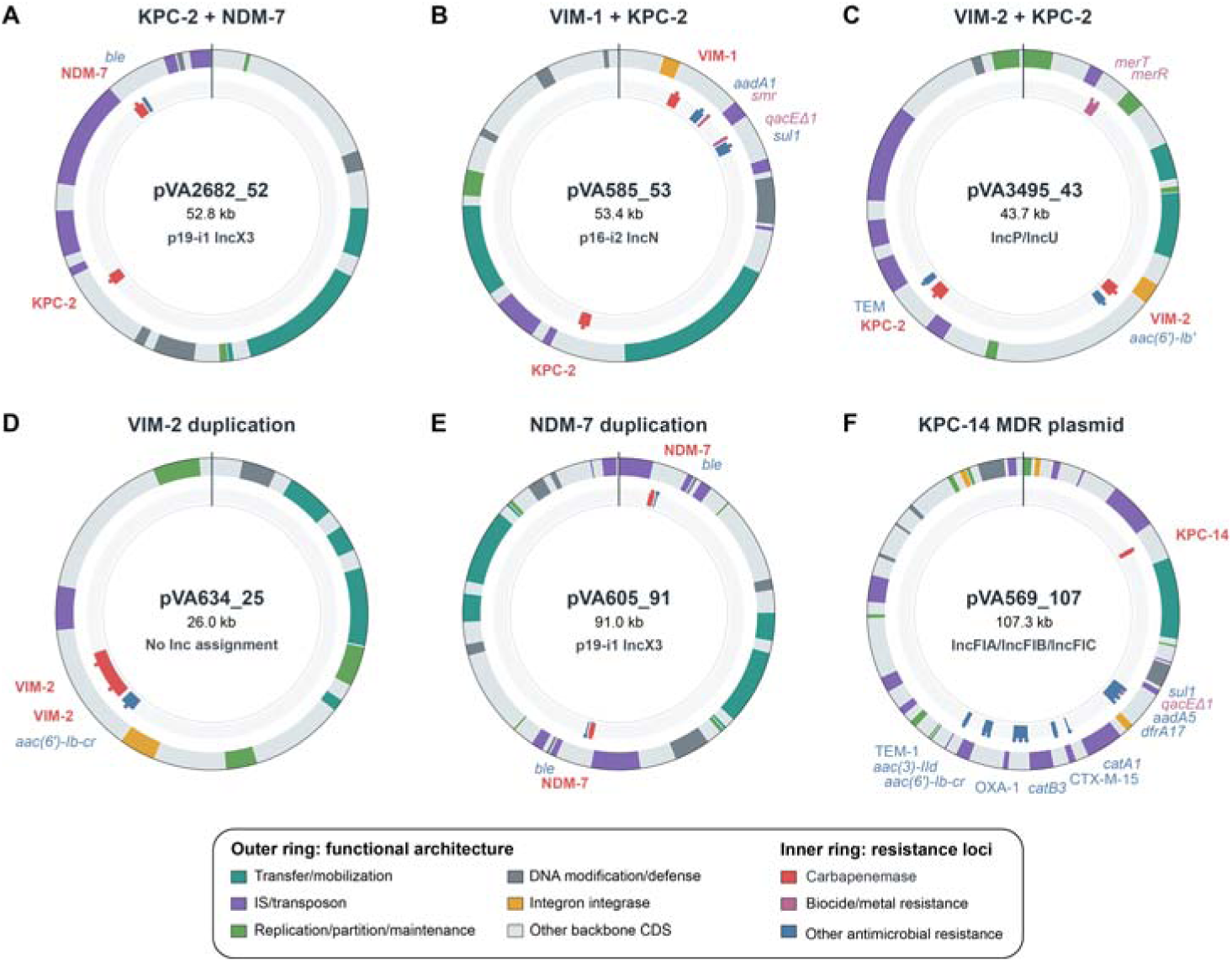
Exceptional architectures of carbapenemase-carrying plasmids. Circular maps show plasmids carrying multiple carbapenemases, duplicated carbapenemase genes, or an uncommon multidrug-resistance configuration. (A) pVA2682_52, a p19-i1 IncX3 plasmid carrying *bla*_KPC-2_ and *bla*_NDM-7_; (B) pVA585_53, a p16-i2 IncN plasmid carrying *bla*_VIM-1_ and *bla*_KPC-2_; (C) pVA3495_43, an IncP/IncU plasmid carrying *bla*_VIM-2_ and *bla*_KPC-2_; (D) pVA634_25, with two copies of *bla*_VIM-2_ and no conventional Inc/Col assignment; (E) pVA605_91, a p19-i1 IncX3 plasmid with two copies of *bla*_NDM-7_; and (F) pVA569_107, an IncFIA/IncFIB/IncFIC multidrug-resistance plasmid carrying the uncommon *bla*_KPC-14_ allele. The outer track classifies coding sequences by functional category, whereas the inner track shows carbapenemases and other antimicrobial-, biocide-, and metal-resistance determinants. Gene labels follow the corresponding resistance-class colors. The radial line at 12 o’clock indicates the sequence origin. Genomic positions are shown relative to each plasmid; circle diameters are not proportional to plasmid size.

Gene duplication provided a second exceptional configuration (Fig. 6D,E). The 26.0-kb plasmid pVA634_25 lacked a conventional Inc/Col assignment and contained two copies of *bla*_VIM-2_, whereas the 91.0-kb IncX3 p19-i1 plasmid pVA605_91 carried two copies of *bla*_NDM-7_. Their unrelated backbones and distinct resistance regions indicate independent duplication or rearrangement events rather than inheritance from a common duplicated architecture.

Singleton plasmids further expanded the resistance repertoire. The 107.3-kb non-mobilizable multireplicon IncF plasmid pVA569_107 carried the uncommon *bla*_KPC-14_ allele together with *bla*_CTX-M-15_, *bla*_OXA-1_, and *bla*_TEM-1_ within a complex multidrug-resistance region (Fig. 6F). In contrast, the conjugative IncC plasmid pVA2817_162 carried *bla*_NDM-1_, an AmpC-type *bla*_CMY_ gene, additional antimicrobial-resistance determinants, and a mercury-resistance region. These examples show how uncommon carbapenemase alleles can occur within larger resistance regions and become linked to additional antibiotic-or metal-resistance determinants.

Together, these plasmids revealed two complementary routes to carbapenemase co-production. Recurrent backbones disseminated carbapenemases across species, while less frequent and structurally exceptional plasmids expanded this repertoire through acquisition of additional resistance cargo, co-localization of carbapenemases, gene duplication, and allelic variation. These patterns point to integrons and transposable regions as a second layer of mobility capable of assembling and redistributing carbapenemase loci across distinct plasmid backbones.

### Conserved integrons and transposon-associated modules underlie the modular distribution of carbapenemase genes

To distinguish circulation of complete plasmids from movement of their resistance modules, we compared the genetic environments of carbapenemase genes across the closed genomes (Fig. 7). Three recurring processes emerged: conservation of integrons across unrelated plasmid backbones, insertion of NDM-associated modules into plasmid and chromosomal compartments, and structural or allelic diversification of KPC-containing regions.

**Figure 7.**
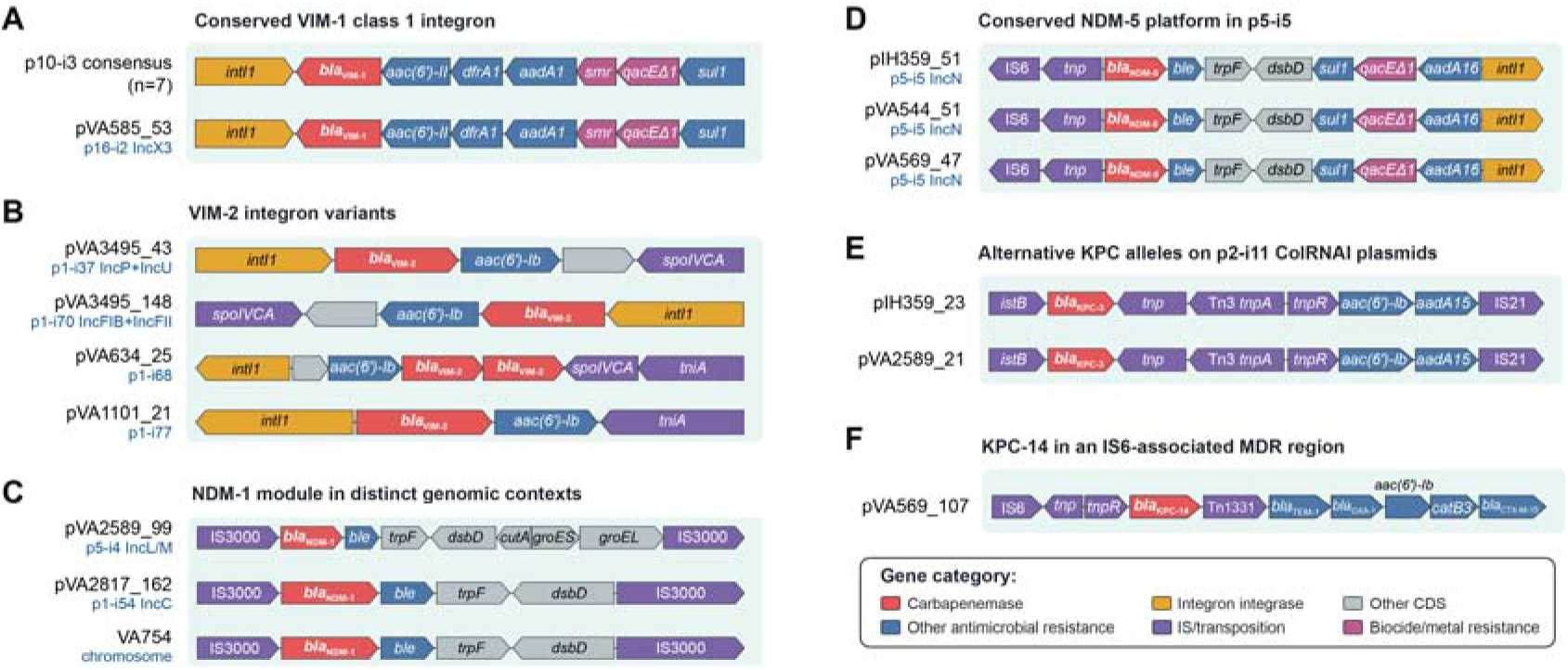
Conserved integrons and transposon-associated modules underlying the modular distribution of carbapenemase genes. (A) Conserved 8,075-bp *bla*_VIM-1_-containing class 1 integron in seven IncHI2A p10-i3 plasmids and in the unrelated IncN plasmid pVA585_53. (B) *bla*_VIM-2_ integron variants in pVA3495_43, pVA3495_148, pVA634_25, and pVA1101_21, including opposite orientations, gene duplication, and shortened configurations. (C) Conserved 11,834-bp IS3000-associated, Tn125-derived *bla*_NDM-1_–*ble* module in the IncL/M-related plasmid pVA2589_99, the IncC plasmid pVA2817_162, and the chromosome of *K. pneumoniae* VA754. (D) Conserved 6,132-bp *bla*_NDM-5_–*ble* platform in the IncN p5-i5 plasmids pIH359_51, pVA544_51, and pVA569_47. (E) Related p2-i11 ColRNAI backbones carrying alternative KPC alleles: *bla*_KPC-3_ in pIH359_23 and *bla*_KPC-66_ in pVA2589_21. (F) IS6-associated multidrug-resistance region carrying *bla*_KPC-14_ in the multireplicon IncFIA/IncFIB/IncFIC plasmid pVA569_107. Arrows indicate coding sequences and their orientation. Colors denote functional categories rather than gene identity; gene names are shown where indicated. Gene lengths and intergenic distances are not drawn to scale.

The eight *bla*_VIM-1_-carrying plasmids provided the clearest evidence of a conserved platform shared by distinct plasmid lineages (Fig. 7A). Seven belonged to p10-i3, whereas pVA585_53 belonged to the unrelated p16-i2 lineage. Despite differences in backbone, size, and bacterial host, all shared an 8,075-bp conserved region containing the same class 1 integron: *intI1*, *bla*_VIM-1_, *aac(6*′*)-Il*, *dfrA1*, *aadA1*, *smr*, *qacE*Δ*1*, and *sul1*. Related VIM-1 integrons have been identified across bacterial genera and plasmid types internationally, supporting an important role for class 1 integrons in disseminating this allele (Matsumura et al., 2017). In Chile, pVA585_53 was previously transferred by conjugation from *K. pneumoniae* ST23 to *E. coli*, and its VIM-1 integron was interpreted as an insertion into an IncN/KPC-2 backbone already circulating in South America (Gálvez-Silva et al., 2024). Its occurrence here in seven IncHI2A-associated plasmids places that event within a broader pattern of integron movement between distinct conjugative backbones. Multiple copies of *sul1* and *qacE*Δ*1* in several p10-i3 plasmids were embedded in larger repeated resistance regions, providing additional evidence of duplication and remodeling of integron-associated modules.

The four *bla*_VIM-2_-containing plasmids displayed greater structural variation (Fig. 7B). Two occurred in *C. portucalensis* VA3495: the IncP/IncU plasmid pVA3495_43 and the unrelated IncFIB/IncFII plasmid pVA3495_148. Their complete *bla*_VIM-2_-containing class 1 integrons were equivalent, indicating capture of the same resistance platform by distinct plasmid backbones. In contrast, pVA634_25 contained a rearranged integron with two copies of *bla*_VIM-2_, whereas pVA1101_21 carried a shortened configuration. These architectures extend a long-standing regional association between *bla*_VIM-2_ and class 1 integrons. VIM-2 was first reported in Latin America in a Chilean *P. aeruginosa* isolate within a class 1 integron, and related platforms have subsequently been found in diverse Gram-negative hosts in South America (Mendes et al., 2004). In the present Enterobacterales, VIM-2 dissemination therefore involved both conservation of an integron across unrelated plasmids and structural remodeling leading to truncation, rearrangement, and gene duplication.

The sole chromosomal carbapenemase, *bla*_NDM-1_ in *K. pneumoniae* VA754, occurred within the same conserved 11,834-bp bla_NDM-1_-ble region identified in the IncL/M-related plasmid pVA2589_99 and the IncC plasmid pVA2817_162 (Fig. 7C). The shared region included *bla*_NDM-1_, *ble*, *trpF*, *dsbD*, and downstream chaperonin genes and was associated with IS3000 elements. This organization resembles Tn3000, an IS3000-bounded element containing a Tn125-derived NDM-1 module initially characterized in Brazil and subsequently detected in geographically and taxonomically diverse Enterobacterales (Campos et al., 2015). A Tn3000-associated *bla*_NDM-1_ region has also been reported on a large conjugative plasmid from a Chilean *K. pneumoniae* isolate (Quezada-Aguiluz et al., 2022). Because the precise boundaries and terminal features of the structures identified here require formal validation, we conservatively describe them as IS3000-associated, Tn125-derived modules. Their occurrence in unrelated plasmids and in the chromosome of VA754 indicates movement across both plasmid-backbone and replicon-compartment boundaries, although the direction of transfer cannot be established.

The three *bla*_NDM-5_-positive p5-i5 plasmids showed a contrasting pattern in which a conserved module circulated with the plasmid backbone (Fig. 7D). Each contained the same 6,132-bp bla_NDM-5_–ble region associated with IS6-and IS91-family elements and *sul1*, resembling the widely distributed NDM-5 module composed of a truncated ISAba125 region, *bla*_NDM-5_, *ble*, *trpF*, and *dsbD* and frequently remodeled by IS26-and IS91-family elements. Its occurrence on three compact IncN plasmids expands the South American plasmid backgrounds associated with NDM-5, previously reported mainly on larger IncF or multireplicon plasmids (Alvarez et al., 2024; González-Espinosa et al., 2024; Magallanes et al., 2025). Conservation across three unrelated *K. pneumoniae* STs supports horizontal dissemination of p5-i5 with limited remodeling.

The closely related p2-i11 plasmids carried *bla*_KPC-3_ and *bla*_KPC-66_ within highly conserved Tn3-and IS21-associated regions (Fig. 7E). Conservation of both the backbone and carbapenemase environment provides structural support for the allelic-diversification scenario discussed above, although independent acquisition cannot be excluded.

The uncommon *bla*_KPC-14_ allele occurred within an approximately 22-kb IS6-rich multidrug-resistance region of pVA569_107, together with a Tn1331-related resolvase and additional β-lactam resistance genes (Fig. 7F). This context differed from the IncN/Tn4401b plasmid in which KPC-14 was initially characterized (Niu et al., 2020) and from later IncFII/IncR plasmids carrying the allele within IS26-associated non-Tn4401 elements (Wang et al., 2023). The Chilean structure therefore represents another genetic background for this uncommon allele. Its boundaries remain unresolved, precluding assignment to a defined transposon. Moreover, although the region contained transfer-associated genes, the complete plasmid was classified as non-mobilizable, suggesting that these functions may be incomplete or disrupted.

These architectures reveal two nested scales of carbapenemase dissemination. Conserved plasmid backbones, exemplified by the NDM-5-positive p5-i5 lineage, circulated among unrelated bacterial clones. At a finer scale, class 1 integrons and insertion-sequence-associated modules connected unrelated plasmids and, for *bla*_NDM-1_, plasmid and chromosomal compartments. Structural remodeling of these modules generated duplicated genes, shortened integrons, and alternative carbapenemase alleles. Complete hybrid assemblies were essential for resolving these relationships and showed that carbapenemase co-production emerged through reassortment at both the plasmid-backbone and nested mobile-element levels.

### Representative isolates reveal distinct within-cell routes to carbapenemase co-production

Genome-resolved analysis showed that similar dual-carbapenemase phenotypes arose through different within-cell arrangements. VA3495, VA605, and VA585 illustrate, respectively, resistance redundancy, convergence of recurrent plasmid lineages, and coexistence of specialized resistance and virulence plasmids (Figs. 4–7; Table S4).

In *C. portucalensis* VA3495, two unrelated plasmids carried equivalent *bla*VIM-2-containing class 1 integrons. One also carried *bla*_KPC-2_, combining two carbapenemases within a single mobilizable plasmid with redundant carriage of the *bla*_VIM-2_ platform on a second conjugative replicon. This arrangement is consistent with transfer or duplication of the integron between co-resident plasmids, although the direction and timing of this event cannot be inferred. It reveals a level of resistance redundancy not captured by describing VA3495 simply as a VIM-2/KPC-2 producer.

A distinct configuration occurred in *E. hormaechei* ST114 isolate VA605. Its IncX3 p19-i1 plasmid pVA605_91 contained two copies of *bla*_NDM-7_, while the co-resident IncHI2A p10-i3 plasmid pVA605_296 carried *bla*_VIM-1_ within the conserved class 1 integron associated with this lineage. VA605 therefore combined two recurrent conjugative plasmid lineages and carried three metallo-β-lactamase gene copies belonging to two enzyme families.

This combination is biologically notable because dual-carbapenemase profiles are not necessarily functionally equivalent. KPC enzymes are class A serine β-lactamases, whereas NDM and VIM are zinc-dependent class B metallo-β-lactamases, resulting in distinct catalytic mechanisms, hydrolytic spectra, and inhibitor profiles (Tooke et al., 2019). KPC/NDM combinations therefore unite mechanistically different enzyme classes, whereas NDM/VIM combinations pair two metallo-β-lactamases with overlapping, although not identical, activities. In VA605, two copies of *bla*_NDM-7_ and one of *bla*_VIM-1_ could increase enzymatic redundancy or total carbapenemase dosage without producing an equivalent expansion of substrate range. This remains a hypothesis because gene expression and enzyme activity were not measured.

The biological consequences of this configuration are also likely to depend on the bacterial and plasmid backgrounds. Copy number can influence β-lactam resistance, and the fitness effects of metallo-β-lactamase production vary with the enzyme, expression level, bacterial host, and zinc availability. NDM enzymes appear comparatively well adapted to expression across Enterobacterales, whereas some VIM enzymes impose measurable fitness or envelope-stress costs under particular conditions (San Millan et al., 2015; López et al., 2019; Cheung et al., 2021; Tu et al., 2025). Interactions between co-resident resistance plasmids can likewise range from additional burden to neutral or beneficial effects (Lee and Ko, 2021). Recovery of VA605 from a clinical infection shows that simultaneous maintenance of the NDM-7 and VIM-1 platforms, including *bla*_NDM-7_ duplication, is possible in an ST114 background. Whether this configuration is stable, costly, or selectively advantageous will require measurements of expression, plasmid stability, and competitive fitness. Its two structurally unrelated conjugative plasmids nevertheless provide separate potential routes for horizontal dissemination. Multiple-carbapenemase isolates should therefore be evaluated according to enzyme class, gene dosage and expression, plasmid mobility, and host-specific fitness effects rather than only by the number of carbapenemase genes detected.

*K. pneumoniae* VA585 provided a previously characterized example of resistance–virulence convergence through specialized co-resident plasmids. Its conjugative IncN plasmid carried *bla*_KPC-2_ and *bla*_VIM-1_, whereas a KpVP-1-like plasmid encoded aerobactin, salmochelin, and *rmp* genes. Previous work established VA585 as a carbapenem-resistant hypervirulent ST23 isolate and demonstrated transfer of its resistance plasmid to *E. coli* (Gálvez-Silva et al., 2024). VA585 therefore illustrates a contrasting route to convergence based on functional specialization between independently maintained resistance and virulence replicons.

Together, these examples show that similar carbapenemase profiles can arise from different genomic arrangements. Distinguishing gene duplication, co-localization on a single plasmid, and carriage on separate transferable replicons is therefore important for understanding how these determinants may persist and disseminate.

## CONCLUSIONS

Multiple-carbapenemase-producing Enterobacterales became increasingly frequent among isolates received through national surveillance in Chile from 2018 to 2024, alongside diversification in bacterial hosts, carbapenemase combinations, and geographic distribution. Phylogenomic analysis of the sequenced subset showed that this pattern was not attributable to a single bacterial clone. Instead, co-production occurred across unrelated species and lineages, reflecting contributions from both clonal persistence and repeated horizontal acquisition.

Complete genomes showed that carbapenemase dissemination involved recurrent conjugative plasmid lineages circulating across bacterial hosts, together with integrons and transposon-associated modules shared among distinct plasmid backbones. Numerous lower-frequency and singleton plasmids, dual-carbapenemase configurations, and gene duplications further revealed a broad and incompletely sampled plasmidome. Gene-level carbapenemase detection alone therefore cannot distinguish dissemination by bacterial clones, complete plasmids, or nested resistance platforms.

Genomic surveillance that resolves complete plasmid backbones and their mobile genetic contexts will be necessary to identify high-risk platforms, track their distribution across bacterial hosts, and monitor the continuing diversification of multiple-carbapenemase-producing Enterobacterales in Chile and the wider region.

## ACKNOWLEDGEMENTS

The authors acknowledge the personnel of the Instituto de Salud Pública de Chile involved in national antimicrobial-resistance surveillance and isolate preservation. Patricio Arros acknowledges support from the Fundación María Ghilardi Venegas fellowship.

## AUTHOR CONTRIBUTIONS

Nicolás Guerra-Leal: Data curation, Formal analysis, Investigation, Methodology, Software, Validation, Visualization, Writing – original draft, Writing – review & editing. Nahuel Pilquinao: Data curation, Formal analysis, Investigation, Methodology, Software, Validation, Visualization, Writing – original draft, Writing – review & editing. Camilo Berríos-Pastén: Formal analysis, Investigation, Methodology, Supervision, Validation. Patricio Arros: Formal analysis, Investigation, Methodology, Supervision, Validation. Matías Gálvez-Silva: Formal analysis, Investigation, Validation. Joaquín Acosta: Formal analysis, Investigation, Validation. Aaron Ascencio: Data curation, Formal analysis, Visualization. Erika Valdivia: Data curation, Formal analysis, Investigation, Visualization. Rosalba Lagos: Project administration, Supervision, Writing – review & editing. Francisco P. Chávez: Project administration, Supervision, Writing – review & editing. Ingrid Araya: Data curation, Investigation, Methodology, Resources, Validation. Rodrigo Iglesias: Data curation, Investigation, Methodology, Resources, Validation. Pamela Araya: Data curation, Project administration, Resources, Supervision, Validation. Juan Carlos Hormazábal: Project administration, Resources, Supervision, Validation. Aura Villamil: Conceptualization, Data curation, Formal analysis, Project administration, Resources, Supervision, Validation, Writing – original draft, Writing – review & editing. Andrés E. Marcoleta: Conceptualization, Funding acquisition, Project administration, Resources, Supervision, Writing – original draft, Writing – review & editing.

## DATA AVAILABILITY

The complete genome assemblies and raw sequencing reads generated in this study are being deposited in the NCBI databases. The BioProject accession and individual assembly and read accession numbers will be added to a revised version of the preprint and provided in Table S3 once assigned. The remaining data supporting the findings of this study are available in the Supplementary Information.

## ETHICS STATEMENT

This study analyzed bacterial isolates and de-identified metadata collected through routine national antimicrobial-resistance surveillance.

## SUPPLEMENTARY FIGURE LEGENDS

**Figure S1. Phylogenomic context of *Klebsiella pneumoniae*.** Maximum-likelihood phylogeny of the eight study isolates and publicly available *K. pneumoniae* genomes, inferred with IQ-TREE from the concatenated core-gene alignment generated with Panaroo and midpoint rooted. Inner annotation rings indicate country of origin and isolation year, while colored sectors identify the sequence types shown in the key. Outer tracks indicate the β-lactamase families listed beside the tree, grouped as extended-spectrum β-lactamases (ESBLs) and carbapenemases (CPs). Red arrowheads identify isolates sequenced in this study, and black arrowheads indicate strains carrying two or more carbapenemase genes. The scale bar represents nucleotide substitutions per site; N/A indicates unavailable metadata.

**Figure S2. Phylogenomic context of *Enterobacter hormaechei*.** Maximum-likelihood phylogeny of the eight study isolates and publicly available *E. hormaechei* genomes, inferred with IQ-TREE from the concatenated core-gene alignment generated with Panaroo and midpoint rooted. Inner annotation rings indicate country of origin and isolation year, while colored sectors identify the sequence types shown in the key. Outer tracks indicate the β-lactamase families listed beside the tree, grouped as extended-spectrum β-lactamases (ESBLs) and carbapenemases (CPs). Red arrowheads identify isolates sequenced in this study, and black arrowheads indicate strains carrying two or more carbapenemase genes. The scale bar represents nucleotide substitutions per site; N/A indicates unavailable metadata.

**Figure S3. Phylogenomic context of *Citrobacter freundii*.** Maximum-likelihood phylogeny of the four study isolates and publicly available *C. freundii* genomes, inferred with IQ-TREE from the concatenated core-gene alignment generated with Panaroo and midpoint rooted. Inner annotation rings indicate country of origin and isolation year, while colored sectors identify the sequence types shown in the key. Outer tracks indicate the β-lactamase families listed beside the tree, grouped as extended-spectrum β-lactamases (ESBLs) and carbapenemases (CPs). Red arrowheads identify isolates sequenced in this study, and black arrowheads indicate strains carrying two or more carbapenemase genes. The scale bar represents nucleotide substitutions per site; N/A indicates unavailable metadata.

**Figure S4. Phylogenomic context of *Citrobacter braakii*.** Maximum-likelihood phylogeny of the study isolate and publicly available *C. braakii* genomes, inferred with IQ-TREE from the concatenated core-gene alignment generated with Panaroo and midpoint rooted. Inner annotation rings indicate country of origin and isolation year, while colored sectors identify the sequence types shown in the key. Outer tracks indicate the β-lactamase families listed beside the tree, grouped as extended-spectrum β-lactamases (ESBLs) and carbapenemases (CPs). The red arrowhead identifies the isolate sequenced in this study, and black arrowheads indicate strains carrying two or more carbapenemase genes. The scale bar represents nucleotide substitutions per site; N/A indicates unavailable metadata.

**Figure S5. Phylogenomic context of *Citrobacter portucalensis*.** Maximum-likelihood phylogeny of the study isolate and publicly available *C. portucalensis* genomes, inferred with IQ-TREE from the concatenated core-gene alignment generated with Panaroo and midpoint rooted. Inner annotation rings indicate country of origin and isolation year, while colored sectors identify the sequence types shown in the key. Outer tracks indicate the β-lactamase families listed beside the tree, grouped as extended-spectrum β-lactamases (ESBLs) and carbapenemases (CPs). The red arrowhead identifies the isolate sequenced in this study, and black arrowheads indicate strains carrying two or more carbapenemase genes. The scale bar represents nucleotide substitutions per site; N/A indicates unavailable metadata.

**Figure S6. Phylogenomic context of *Enterobacter asburiae*.** Maximum-likelihood phylogeny of the study isolate and publicly available *E. asburiae* genomes, inferred with IQ-TREE from the concatenated core-gene alignment generated with Panaroo and midpoint rooted. Inner annotation rings indicate country of origin and isolation year, while colored sectors identify the sequence types shown in the key. Outer tracks indicate the β-lactamase families listed beside the tree, grouped as extended-spectrum β-lactamases (ESBLs) and carbapenemases (CPs). The red arrowhead identifies the isolate sequenced in this study, and black arrowheads indicate strains carrying two or more carbapenemase genes. The scale bar represents nucleotide substitutions per site; N/A indicates unavailable metadata.

**Figure S7. Phylogenomic context of *Klebsiella aerogenes*.** Maximum-likelihood phylogeny of the two study isolates and publicly available *K. aerogenes* genomes, inferred with IQ-TREE from the concatenated core-gene alignment generated with Panaroo and midpoint rooted. Inner annotation rings indicate country of origin and isolation year, while colored sectors identify the sequence types shown in the key. Outer tracks indicate the β-lactamase families listed beside the tree, grouped as extended-spectrum β-lactamases (ESBLs) and carbapenemases (CPs). Red arrowheads identify isolates sequenced in this study, and black arrowheads indicate strains carrying two or more carbapenemase genes. The scale bar represents nucleotide substitutions per site; N/A indicates unavailable metadata.

**Figure S8. Phylogenomic context of *Klebsiella michiganensis*.** Maximum-likelihood phylogeny of the two study isolates and publicly available *K. michiganensis* genomes, inferred with IQ-TREE from the concatenated core-gene alignment generated with Panaroo and midpoint rooted. Inner annotation rings indicate country of origin and isolation year, while colored sectors identify the sequence types shown in the key. Outer tracks indicate the β-lactamase families listed beside the tree, grouped as extended-spectrum β-lactamases (ESBLs) and carbapenemases (CPs). Red arrowheads identify isolates sequenced in this study, and black arrowheads indicate strains carrying two or more carbapenemase genes. The scale bar represents nucleotide substitutions per site; N/A indicates unavailable metadata.

**Figure S9. Phylogenomic context of *Klebsiella planticola*.** Maximum-likelihood phylogeny of the study isolate and publicly available *K. planticola* genomes, inferred with IQ-TREE from the concatenated core-gene alignment generated with Panaroo and midpoint rooted. Inner annotation rings indicate country of origin and isolation year. Outer tracks indicate the β-lactamase families listed beside the tree, grouped as extended-spectrum β-lactamases (ESBLs) and carbapenemases (CPs). The red arrowhead identifies the isolate sequenced in this study, and black arrowheads indicate strains carrying two or more carbapenemase genes. The scale bar represents nucleotide substitutions per site; N/A indicates unavailable metadata.

**Figure S10. Phylogenomic context of *Leclercia spp.*** Maximum-likelihood phylogeny of the *Leclercia tamurae* study isolate and publicly available *Leclercia* genomes, inferred with IQ-TREE from the concatenated core-gene alignment generated with Panaroo and midpoint rooted. Inner annotation rings indicate country of origin and isolation year. Outer tracks indicate the β-lactamase families listed beside the tree, grouped as extended-spectrum β-lactamases (ESBLs) and carbapenemases (CPs). The red arrowhead identifies the isolate sequenced in this study, and black arrowheads indicate strains carrying two or more carbapenemase genes. The scale bar represents nucleotide substitutions per site; N/A indicates unavailable metadata.

**Figure S11. Phylogenomic context of *Serratia marcescens*.** Maximum-likelihood phylogeny of the study isolate and publicly available *S. marcescens* genomes, inferred with IQ-TREE from the concatenated core-gene alignment generated with Panaroo and midpoint rooted. Inner annotation rings indicate country of origin and isolation year. Outer tracks indicate the β-lactamase families listed beside the tree, grouped as extended-spectrum β-lactamases (ESBLs) and carbapenemases (CPs). The red arrowhead identifies the isolate sequenced in this study, and black arrowheads indicate strains carrying two or more carbapenemase genes. The scale bar represents nucleotide substitutions per site; N/A indicates unavailable metadata.

**Figure S12. Sample-based rarefaction and occupancy of plasmid lineages.** (A) Incidence-based accumulation curve for operational plasmid lineages across the 30 sequenced isolates. The solid line represents analytical rarefaction, the dashed line represents Chao2-based extrapolation, and shaded areas indicate 95% confidence intervals. The horizontal dotted line indicates the bias-corrected Chao2 richness estimate. (B) Distribution of plasmid lineages according to the number of isolates in which they were detected. Operational lineages were defined by exact primary-secondary MOB-cluster pairs. Two plasmids lacking either MOB-cluster assignment were excluded from the analysis.

**Figure S13. Pan-plasmidome structure of the three recurrent plasmid lineages.** Gene-family presence–absence matrices for plasmids assigned to (A) p19-i1 IncX3 carrying *bla*_NDM-7_ (n = 19; 40.9–91.0 kb), (B) p16-i2 IncN carrying *bla*_KPC-2_ (n = 16; 41.3–61.2 kb), and (C) p10-i3 IncHI2A (n = 10; 265.9–375.6 kb) frequently carrying *bla*_VIM-1_. Rows correspond to individual plasmids and columns to gene families identified in the lineage-specific pan-plasmidome analysis; gray cells indicate gene-family presence and white cells indicate absence. Carbapenemase genes and selected accessory regions are indicated, including *ars* and *mer* determinants, integron-associated antimicrobial-resistance genes, and variable regions involved in conjugation, replication, and plasmid maintenance. Gene-family matrices illustrate the conserved backbones and variable accessory regions within each recurrent lineage.

## SUPPLEMENTARY TABLE LEGENDS

**Table S1. Epidemiological, taxonomic, and carbapenemase characteristics of multiple-carbapenemase-producing Enterobacterales identified through Chilean national surveillance, 2018–2024.** One row is provided for each unique ISP isolate record. Ye*ars* correspond to the ISP record identifier; taxonomic assignments and regions reflect surveillance metadata.

**Table S2. Antimicrobial susceptibility profiles of the 30 sequenced study isolates.** Minimum inhibitory concentrations are reported in µg/mL unless a zone diameter in millimeters is specified. Interpretation codes and antimicrobial abbreviations are defined in the table notes.

**Table S3. Sequencing, assembly, and genome-quality metrics for the 30 study isolates.** The table includes genome-based taxonomy, sequence type, completeness and contamination estimates, assembly metrics, plasmid counts, carbapenemases, and extended-spectrum β-lactamases.

**Table S4. Genomic characteristics, mobility, replicon content,** β**-lactamase cargo, and sequence-based classification of the 149 complete plasmids recovered from the 30 study isolates.** One row is provided per plasmid, including size, bacterial host, sequence type and year, carbapenemase copy number, β-lactamase content, conventional Inc/Col replicon assignment, predicted mobility, primary and secondary MOB-cluster identifiers, and study-specific plasmid-lineage designation (pA-iB).

