## Supplementary Figures for "Cross-species dissemination of diverse plasmid lineages and mobile resistance modules shapes carbapenemase co-production in Chilean Enterobacterales"

***Klebsiella pneumoniae***

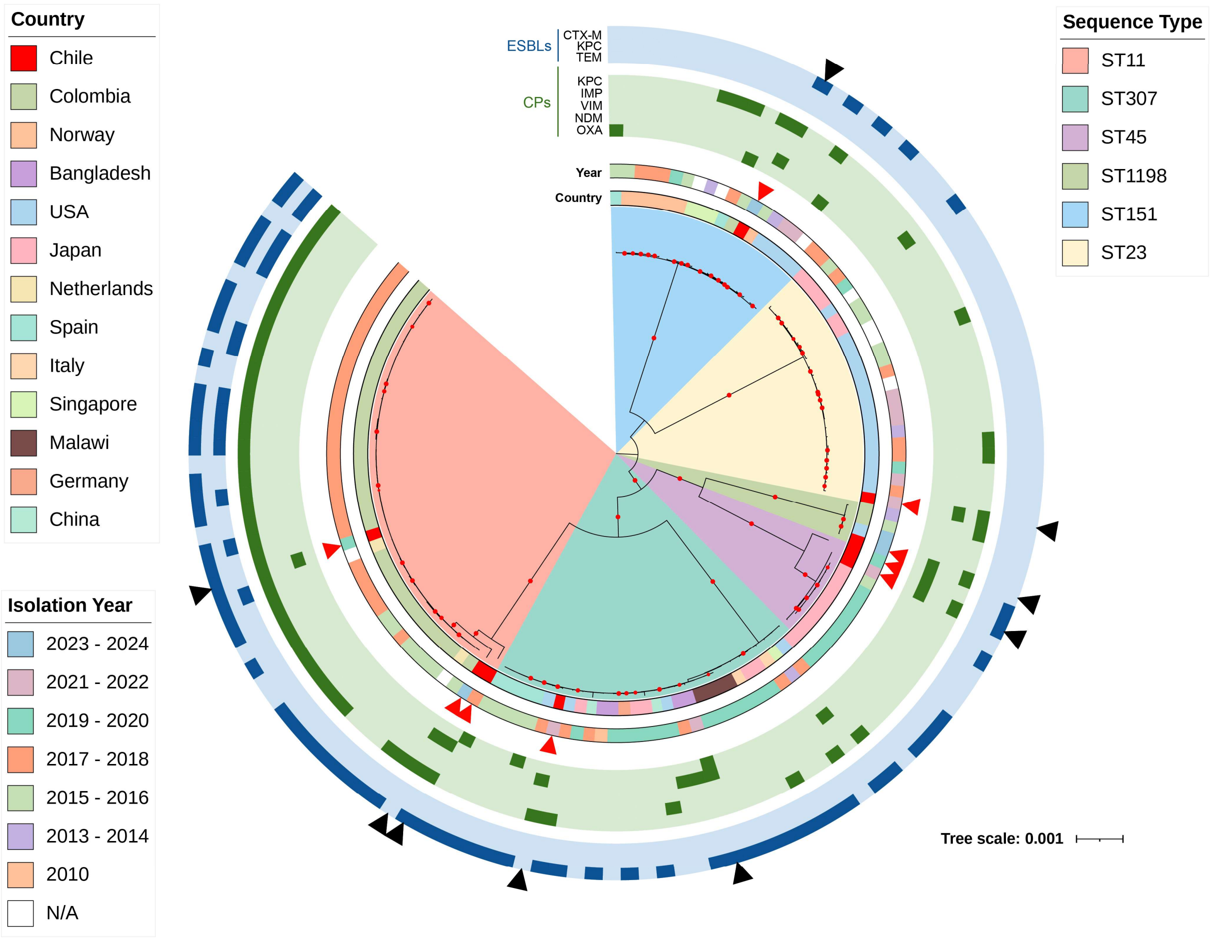


**Figure S1. Phylogenomic context of *Klebsiella pneumoniae*.** Maximum-likelihood phylogeny of the eight study isolates and publicly available *K. pneumoniae* genomes, inferred with IQ-TREE from the concatenated core-gene alignment generated with Panaroo and midpoint rooted. Inner annotation rings indicate country of origin and isolation year, while colored sectors identify the sequence types shown in the key. Outer tracks indicate the β-lactamase families listed beside the tree, grouped as extended-spectrum β-lactamases (ESBLs) and carbapenemases (CPs). Red arrowheads identify isolates sequenced in this study, and black arrowheads indicate strains carrying two or more carbapenemase genes. The scale bar represents nucleotide substitutions per site; N/A indicates unavailable metadata.

***Enterobacter hormaechei***

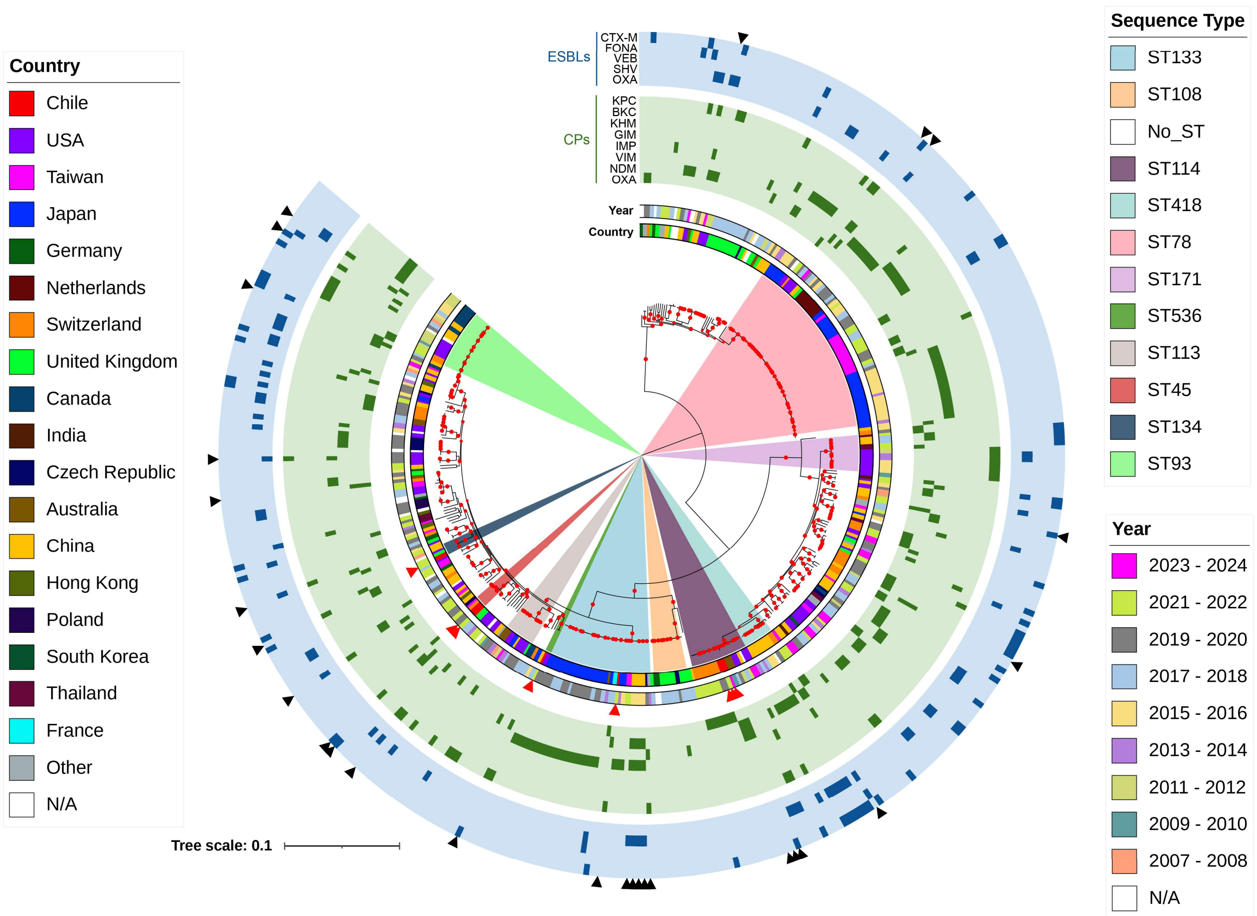


**Figure S2. Phylogenomic context of *Enterobacter hormaechei*.** Maximum-likelihood phylogeny of the eight study isolates and publicly available *E. hormaechei* genomes, inferred with IQ-TREE from the concatenated core-gene alignment generated with Panaroo and midpoint rooted. Inner annotation rings indicate country of origin and isolation year, while colored sectors identify the sequence types shown in the key. Outer tracks indicate the β-lactamase families listed beside the tree, grouped as extended-spectrum β-lactamases (ESBLs) and carbapenemases (CPs). Red arrowheads identify isolates sequenced in this study, and black arrowheads indicate strains carrying two or more carbapenemase genes. The scale bar represents nucleotide substitutions per site; N/A indicates unavailable metadata.

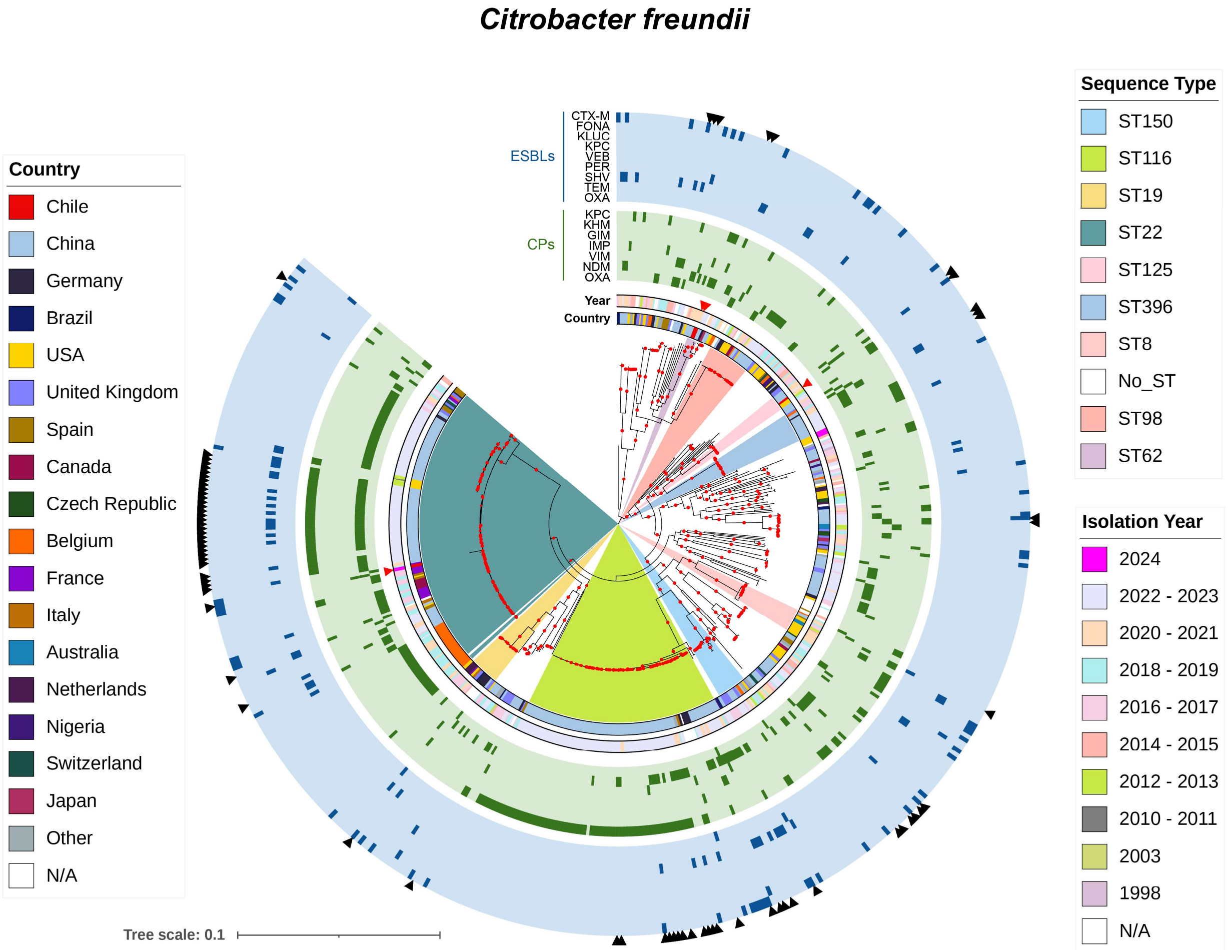


**Figure S3. Phylogenomic context of *Citrobacter freundii*.** Maximum-likelihood phylogeny of the four study isolates and publicly available *C. freundii* genomes, inferred with IQ-TREE from the concatenated core-gene alignment generated with Panaroo and midpoint rooted. Inner annotation rings indicate country of origin and isolation year, while colored sectors identify the sequence types shown in the key. Outer tracks indicate the β-lactamase families listed beside the tree, grouped as extended-spectrum β-lactamases (ESBLs) and carbapenemases (CPs). Red arrowheads identify isolates sequenced in this study, and black arrowheads indicate strains carrying two or more carbapenemase genes. The scale bar represents nucleotide substitutions per site; N/A indicates unavailable metadata.

#### *Citrobacter braakii*

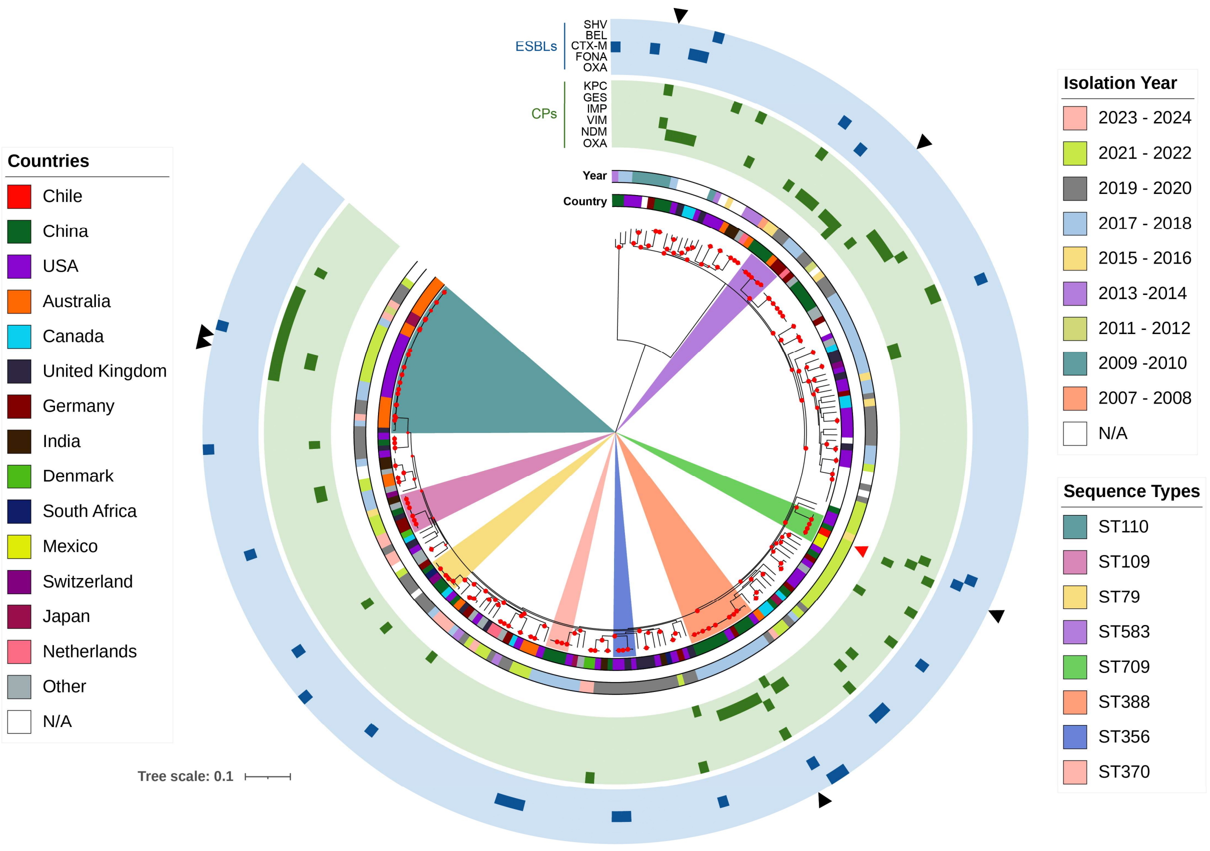


**Figure S4. Phylogenomic context of *Citrobacter braakii*.** Maximum-likelihood phylogeny of the study isolate and publicly available *C. braakii* genomes, inferred with IQ-TREE from the concatenated core-gene alignment generated with Panaroo and midpoint rooted. Inner annotation rings indicate country of origin and isolation year, while colored sectors identify the sequence types shown in the key. Outer tracks indicate the β-lactamase families listed beside the tree, grouped as extended-spectrum β-lactamases (ESBLs) and carbapenemases (CPs). The red arrowhead identifies the isolate sequenced in this study, and black arrowheads indicate strains carrying two or more carbapenemase genes. The scale bar represents nucleotide substitutions per site; N/A indicates unavailable metadata.

*Citrobacter portucalensis*

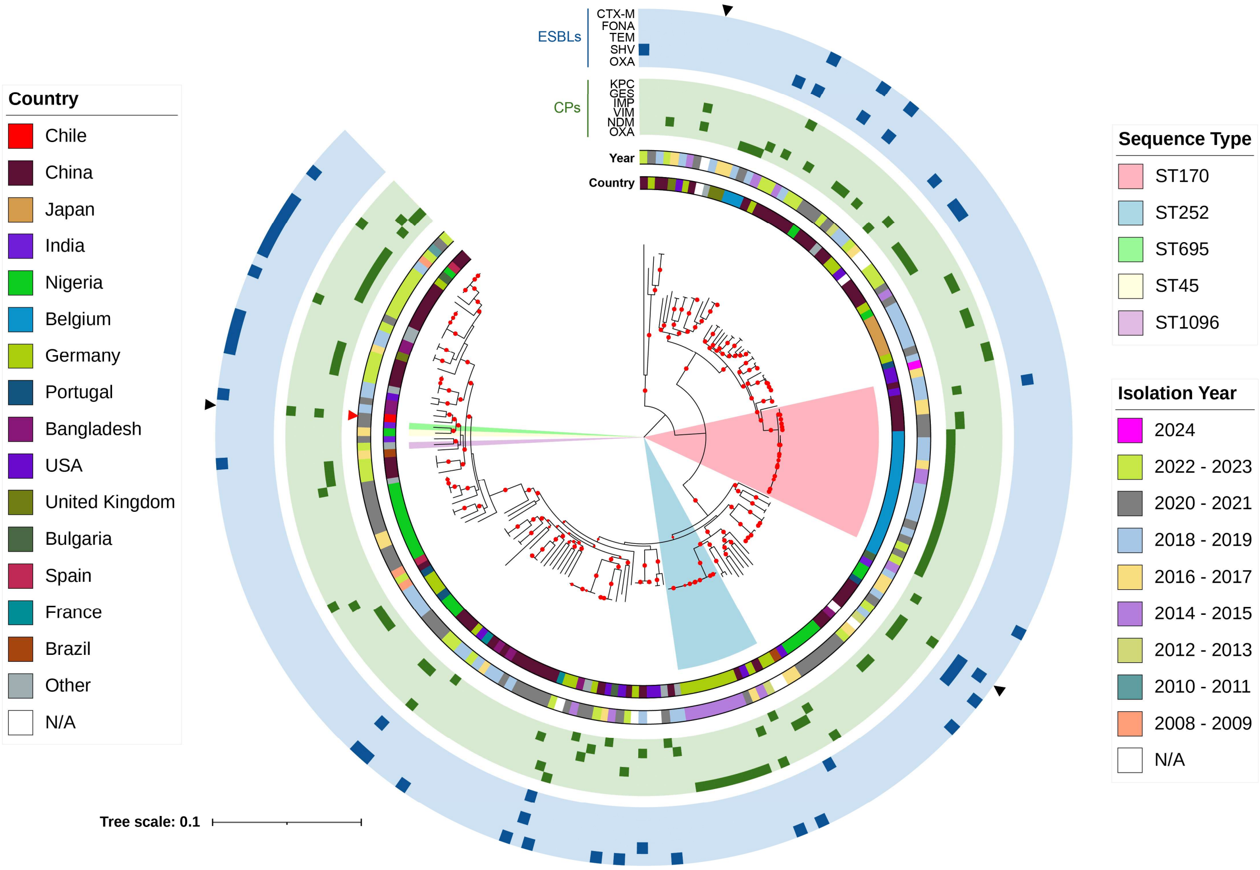


**Figure S5. Phylogenomic context of *Citrobacter portucalensis*.** Maximum-likelihood phylogeny of the study isolate and publicly available *C. portucalensis* genomes, inferred with IQ-TREE from the concatenated core-gene alignment generated with Panaroo and midpoint rooted. Inner annotation rings indicate country of origin and isolation year, while colored sectors identify the sequence types shown in the key. Outer tracks indicate the β-lactamase families listed beside the tree, grouped as extended-spectrum β-lactamases (ESBLs) and carbapenemases (CPs). The red arrowhead identifies the isolate sequenced in this study, and black arrowheads indicate strains carrying two or more carbapenemase genes. The scale bar represents nucleotide substitutions per site; N/A indicates unavailable metadata.

#### *Enterobacter asburiae*

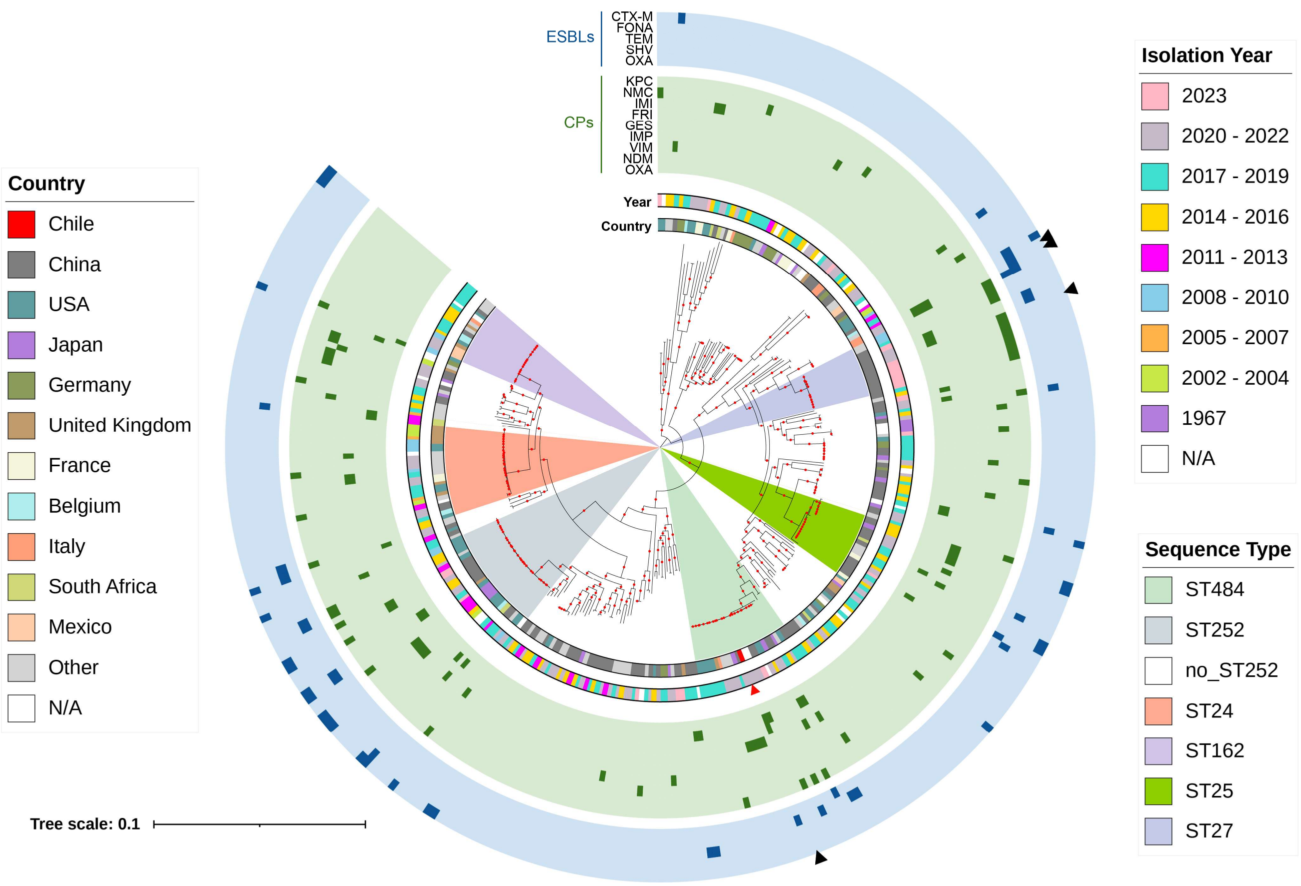


**Figure S6. Phylogenomic context of *Enterobacter asburiae*.** Maximum-likelihood phylogeny of the study isolate and publicly available *E. asburiae* genomes, inferred with IQ-TREE from the concatenated core-gene alignment generated with Panaroo and midpoint rooted. Inner annotation rings indicate country of origin and isolation year, while colored sectors identify the sequence types shown in the key. Outer tracks indicate the β-lactamase families listed beside the tree, grouped as extended-spectrum β-lactamases (ESBLs) and carbapenemases (CPs). The red arrowhead identifies the isolate sequenced in this study, and black arrowheads indicate strains carrying two or more carbapenemase genes. The scale bar represents nucleotide substitutions per site; N/A indicates unavailable metadata.

#### *Klebsiella aerogenes*

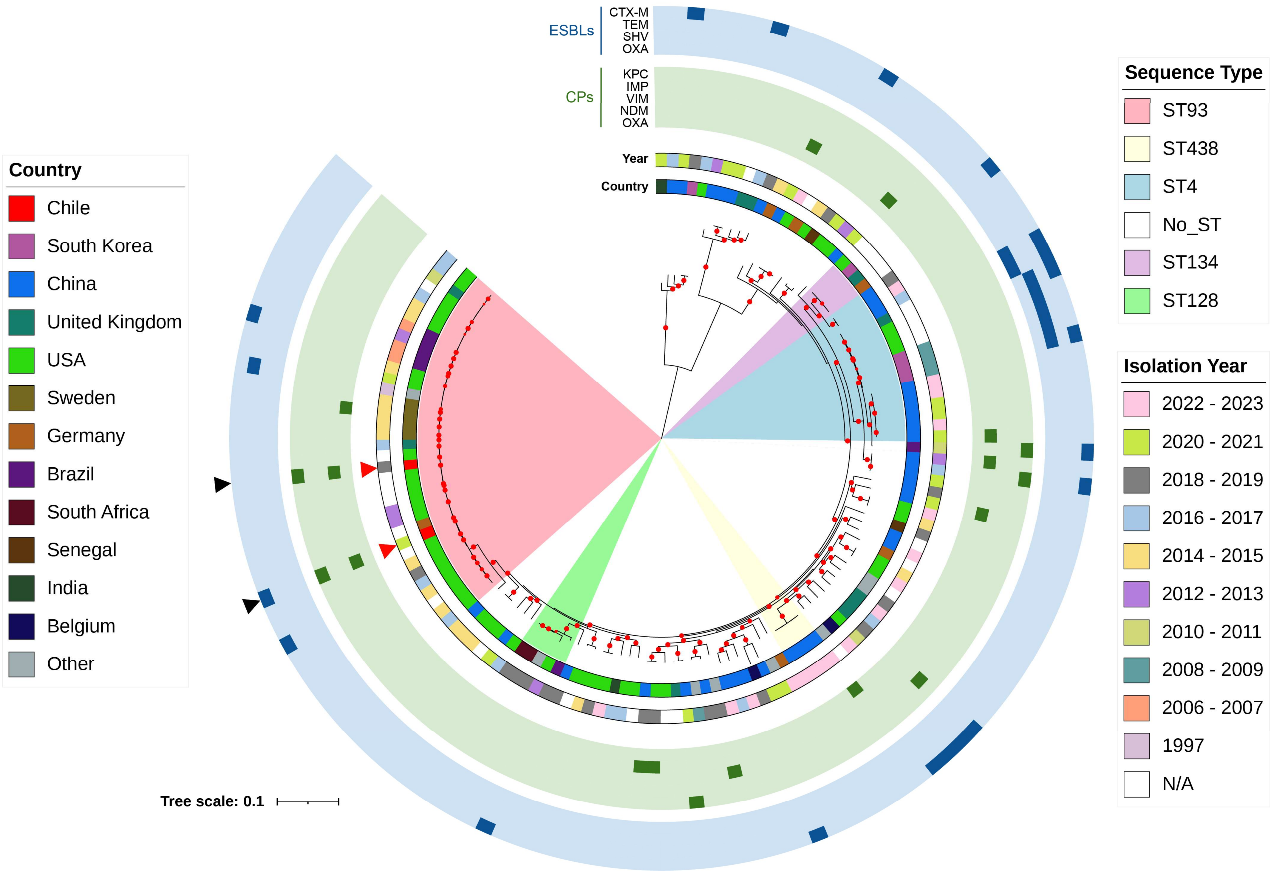


**Figure S7. Phylogenomic context of *Klebsiella aerogenes*.** Maximum-likelihood phylogeny of the two study isolates and publicly available *K. aerogenes* genomes, inferred with IQ-TREE from the concatenated core-gene alignment generated with Panaroo and midpoint rooted. Inner annotation rings indicate country of origin and isolation year, while colored sectors identify the sequence types shown in the key. Outer tracks indicate the β-lactamase families listed beside the tree, grouped as extended-spectrum β-lactamases (ESBLs) and carbapenemases (CPs). Red arrowheads identify isolates sequenced in this study, and black arrowheads indicate strains carrying two or more carbapenemase genes. The scale bar represents nucleotide substitutions per site; N/A indicates unavailable metadata.

### *Klebsiella michiganensis*

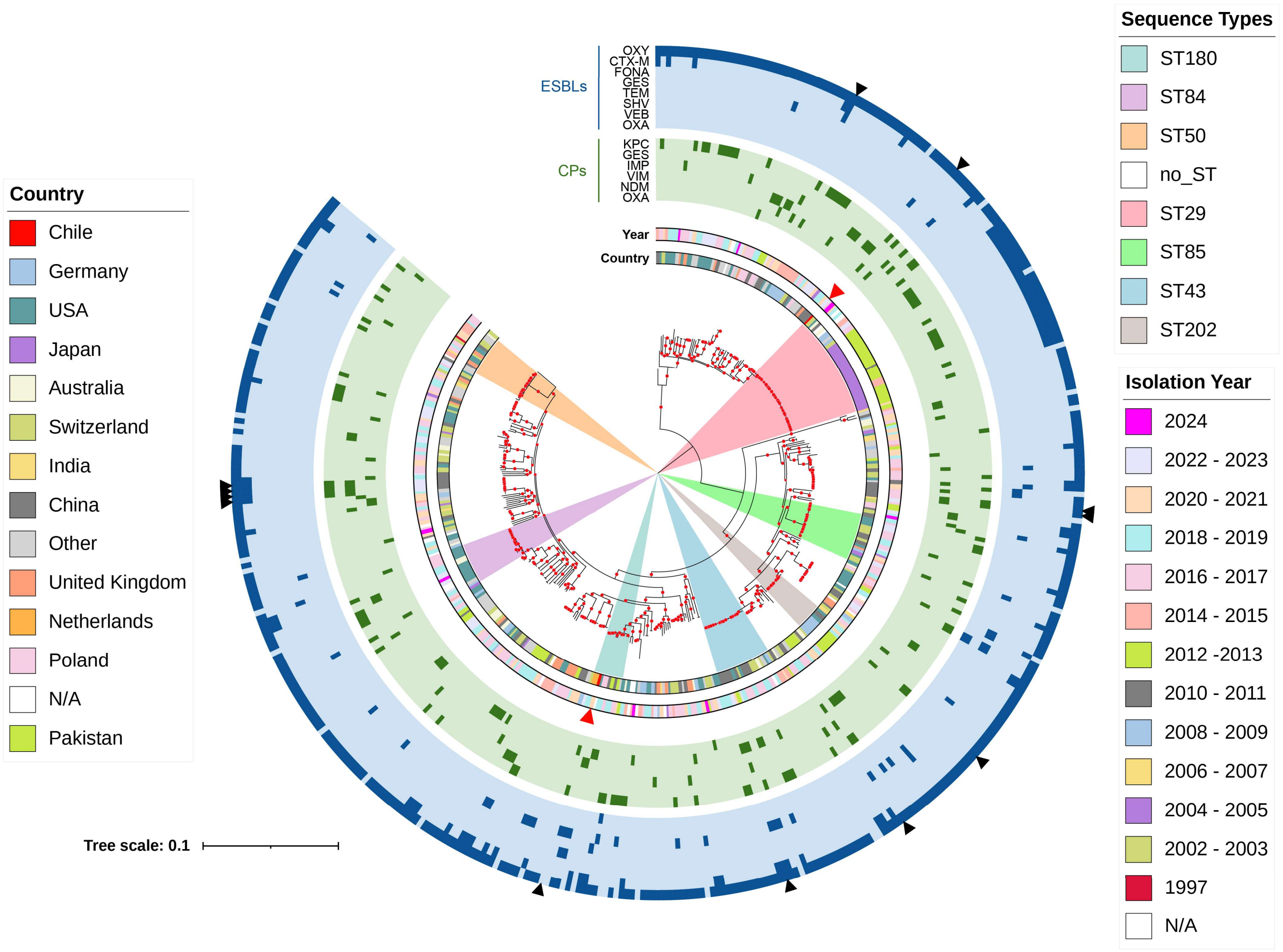


**Figure S8. Phylogenomic context of *Klebsiella michiganensis*.** Maximum-likelihood phylogeny of the two study isolates and publicly available *K. michiganensis* genomes, inferred with IQ-TREE from the concatenated core-gene alignment generated with Panaroo and midpoint rooted. Inner annotation rings indicate country of origin and isolation year, while colored sectors identify the sequence types shown in the key. Outer tracks indicate the β-lactamase families listed beside the tree, grouped as extended-spectrum β-lactamases (ESBLs) and carbapenemases (CPs). Red arrowheads identify isolates sequenced in this study, and black arrowheads indicate strains carrying two or more carbapenemase genes. The scale bar represents nucleotide substitutions per site; N/A indicates unavailable metadata.

#### *Klebsiella planticola*

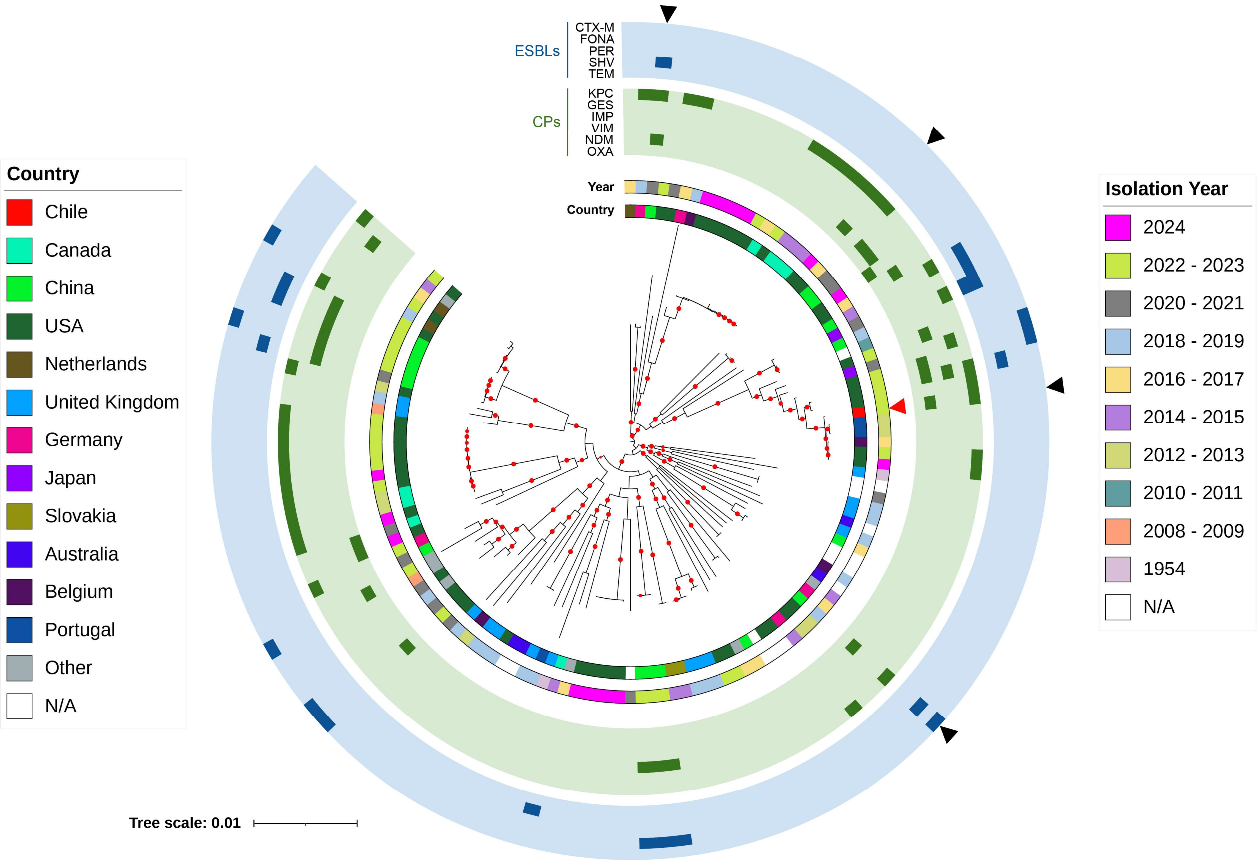


**Figure S9. Phylogenomic context of *Klebsiella planticola*.** Maximum-likelihood phylogeny of the study isolate and publicly available *K. planticola* genomes, inferred with IQ-TREE from the concatenated core-gene alignment generated with Panaroo and midpoint rooted. Inner annotation rings indicate country of origin and isolation year. Outer tracks indicate the β-lactamase families listed beside the tree, grouped as extended-spectrum β-lactamases (ESBLs) and carbapenemases (CPs). The red arrowhead identifies the isolate sequenced in this study, and black arrowheads indicate strains carrying two or more carbapenemase genes. The scale bar represents nucleotide substitutions per site; N/A indicates unavailable metadata.

#### *Leclercia spp.*

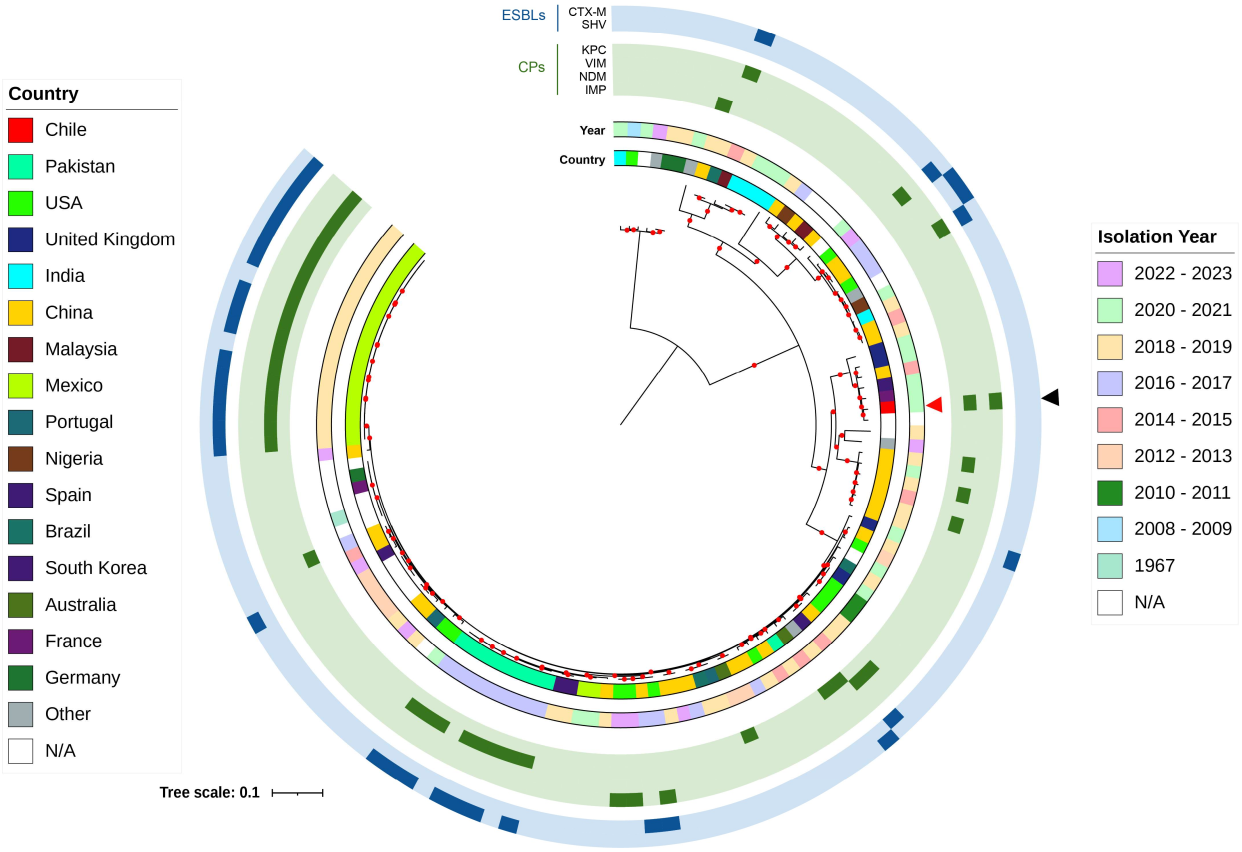


**Figure S10. Phylogenomic context of *Leclercia* spp.** Maximum-likelihood phylogeny of the *Leclercia tamurae* study isolate and publicly available *Leclercia* genomes, inferred with IQ-TREE from the concatenated core-gene alignment generated with Panaroo and midpoint rooted. Inner annotation rings indicate country of origin and isolation year. Outer tracks indicate the β-lactamase families listed beside the tree, grouped as extended-spectrum β-lactamases (ESBLs) and carbapenemases (CPs). The red arrowhead identifies the isolate sequenced in this study, and black arrowheads indicate strains carrying two or more carbapenemase genes. The scale bar represents nucleotide substitutions per site; N/A indicates unavailable metadata.

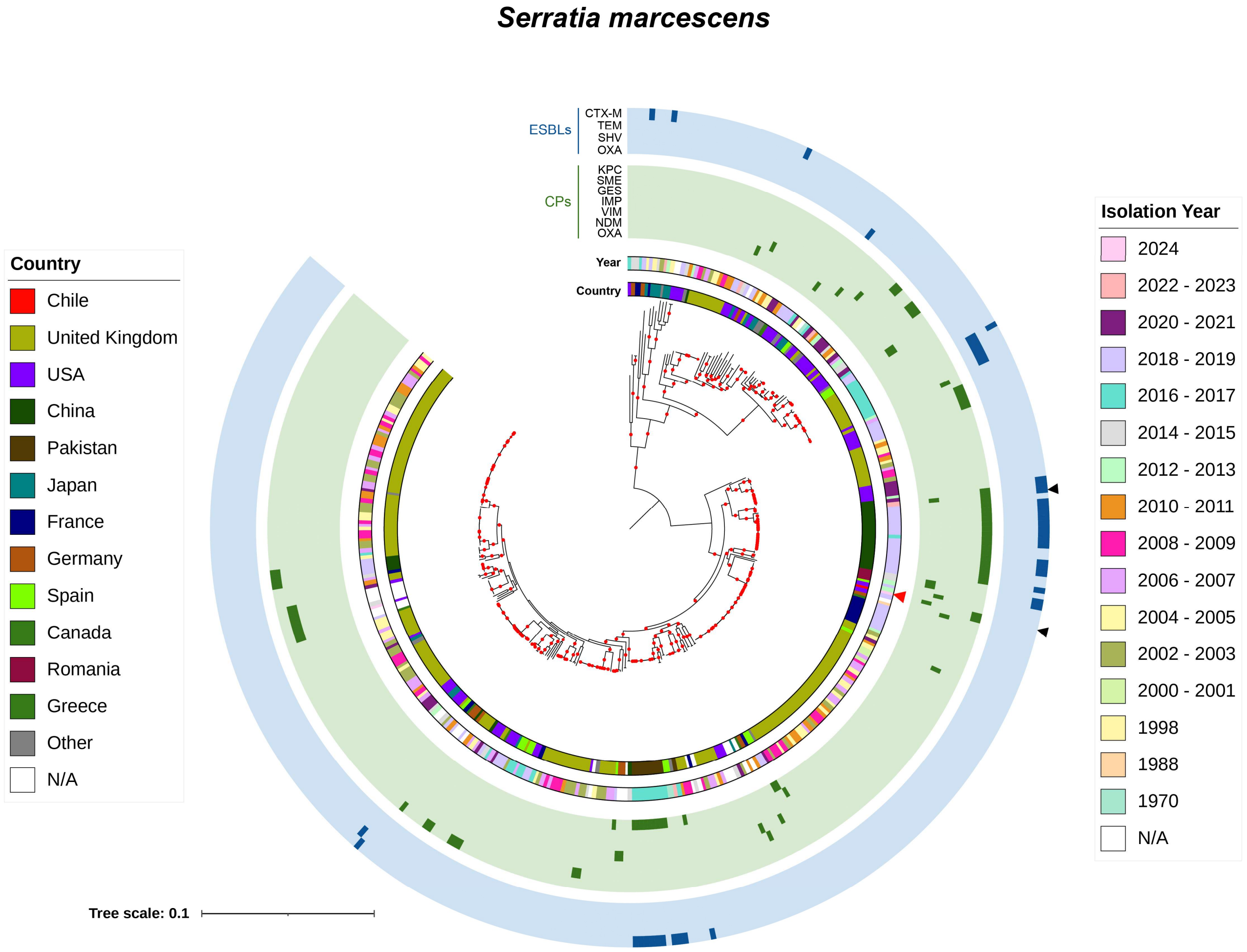


**Figure S11. Phylogenomic context of *Serratia marcescens*.** Maximum-likelihood phylogeny of the study isolate and publicly available *S. marcescens* genomes, inferred with IQ-TREE from the concatenated core-gene alignment generated with Panaroo and midpoint rooted. Inner annotation rings indicate country of origin and isolation year. Outer tracks indicate the β-lactamase families listed beside the tree, grouped as extended-spectrum β-lactamases (ESBLs) and carbapenemases (CPs). The red arrowhead identifies the isolate sequenced in this study, and black arrowheads indicate strains carrying two or more carbapenemase genes. The scale bar represents nucleotide substitutions per site; N/A indicates unavailable metadata.

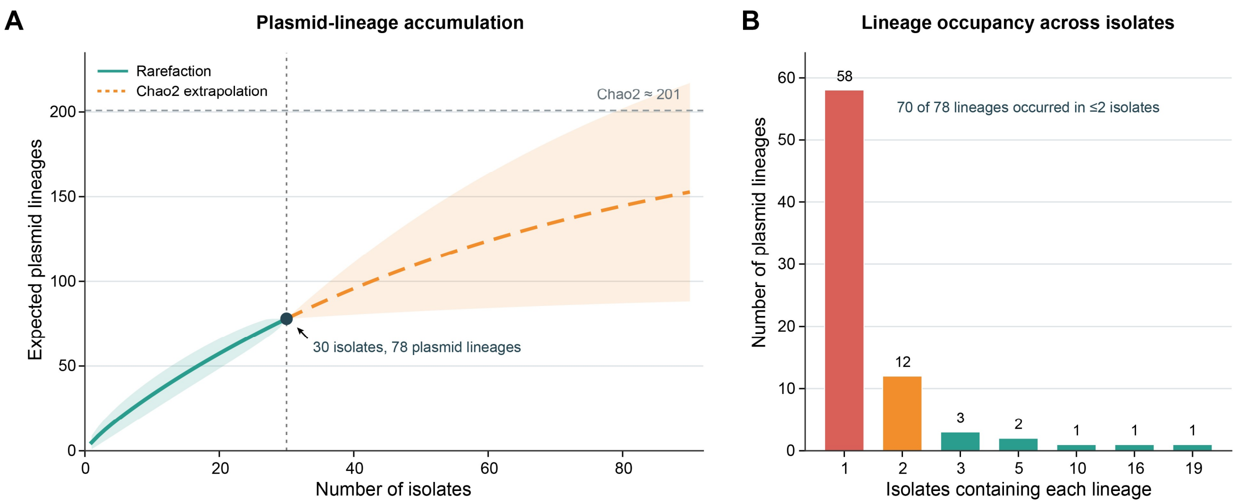


**Figure S12. Sample-based rarefaction and occupancy of plasmid lineages.** (A) Incidence-based accumulation curve for operational plasmid lineages across the 30 sequenced isolates. The solid line represents analytical rarefaction, the dashed line represents Chao2-based extrapolation, and shaded areas indicate 95% confidence intervals. The horizontal dotted line indicates the bias-corrected Chao2 richness estimate. (B) Distribution of plasmid lineages according to the number of isolates in which they were detected. Operational lineages were defined by exact primary-secondary MOB-cluster pairs. Two plasmids lacking either MOB-cluster assignment were excluded from the analysis.

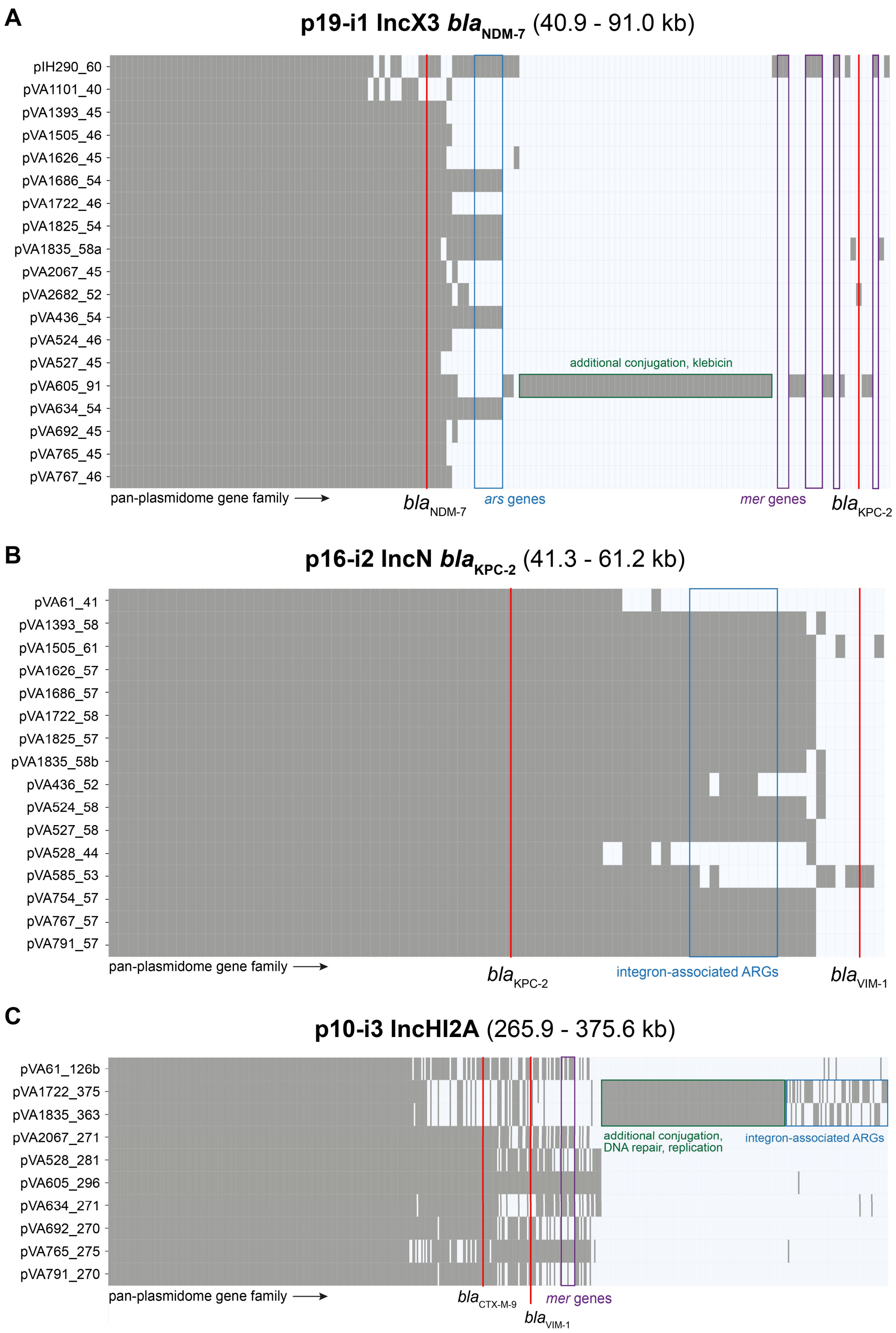


**Figure S13. Pan-plasmidome structure of the three recurrent plasmid lineages.** Gene-family presence–absence matrices for plasmids assigned to (A) p19-i1 IncX3 carrying *bla*$_{NDM-7}$ (n = 19; 40.9–91.0 kb), (B) p16-i2 IncN carrying *bla*$_{KPC-2}$ (n = 16; 41.3–61.2 kb), and (C) p10-i3 IncHI2A (n = 10; 265.9–375.6 kb) frequently carrying *bla*$_{VIM-1}$. Rows correspond to individual plasmids and columns to gene families identified in the lineage-specific pan-plasmidome analysis; gray cells indicate gene-family presence and white cells indicate absence. Carbapenemase genes and selected accessory regions are indicated, including *ars* and *mer* determinants, integron-associated antimicrobial resistance genes, and variable regions involved in conjugation, replication, and plasmid maintenance. Gene-family matrices illustrate the conserved backbones and variable accessory regions within each recurrent lineage.
